# American black bear (*Ursus americanus*) as a potential host for *Campylobacter jejuni*

**DOI:** 10.1101/2025.04.14.648838

**Authors:** Craig T. Parker, Sophia Kathariou, William G. Miller, Steven Huynh, Ben Pascoe, Kerry K. Cooper

## Abstract

The Gram-negative bacterium *Campylobacter jejuni* is part of the commensal gut microbiota of numerous animal species and a leading cause of bacterial foodborne illness in humans. Most complete genomes of *C. jejuni* are from strains isolated from human clinical, poultry, and ruminant samples. Here, we characterized and compared the genomes of *C. jejuni* that were isolated from American black bears in three states in the southeastern United States. Despite the limited sample size (n=9), the isolates displayed substantial genotypic variability, including eight distinct sequence types (STs) and variable gene content encoding surface glycan structures such as capsular polysaccharides (CPS) and lipooligosaccharides (LOS). Phylogenetic analysis identified several *C. jejuni* host generalist strains among the isolates from bears that clustered with isolates from domestic poultry, cattle, and environmental sources. Three isolates (SKBC94, SKBC3, SKBC5) clustered with wildlife-associated strains, exhibiting mutations or deletions in loci associated with cytolethal distending toxin production and oxidative stress resistance, potentially influencing host-specific colonization. Additionally, strains SKBC3 and SKBC5 harbored distinct Entner-Doudoroff (E-D) loci, suggesting a potential evolutionary fitness advantage. This study provides the first evidence of *C. jejuni* colonization in American black bears, highlighting their potential role as reservoirs for diverse *C. jejuni* lineages from both anthropogenic and environmental sources. Further research is needed to determine the prevalence and host specificity of *C. jejuni* strains in black bears and their potential implications for public and wildlife health.

## Introduction

*Campylobacter jejuni* is the leading cause of bacterial gastroenteritis worldwide (1–3) and has adapted to the gastrointestinal tract of various avian and mammalian hosts. Human infections with *C. jejuni* are usually the result of consumption and/or handling of contaminated poultry products, while raw milk and untreated water are also common sources of infection (3–9). Characterization of strain level variation, using methods such as multilocus sequence typing (MLST), has identified genotypes of *C. jejuni* strains found in multiple host species (e.g. domesticated poultry, ruminants, etc.) termed host generalists and genotypes of strains isolated primarily from single reservoir species (e.g. domestic cattle or Guinea pig) termed host specialists (10–14). The genotypes of *C. jejuni* that are seldom associated with human disease have been identified from non-food production animals (14–19). In most cases, these genotypes, such as those *C. jejuni* strains isolated from raccoons and many wild bird species are dissimilar genotypes of strains that cause human disease or colonize livestock animals. These genotypic differences represent signals of host adaptation that are used in public health studies to attribute human infection cases to likely reservoir source(s) (10, 19–22). However, in more urban settings, strains possessing host generalist genotypes can be isolated from multiple hosts species that share the same niche, which complicates source attribution in these cases (12, 16, 18, 19, 23–27).

Occasionally, direct transmission of *C. jejuni* to humans has been linked to wildlife (28–30), and population level analyses of human clinical campylobacteriosis cases attributes a small proportion of infections to strains with wildlife-associated genotypes (12, 18, 19, 31). Shedding of *C. jejuni* in animal feces, especially around water, provides a means for the pathogen to enter new ecological niches, exchange between hosts, and cause infections in humans (18, 32, 33). In the agricultural environment, the proximity of domestic and wild animals allows certain *C. jejuni* lineages to be shared among different hosts (12, 18). Indeed, raccoons and guinea pigs harbor *C. jejuni* strains with both their own host-adapted and also domestic animal-associated genotypes (18, 26).

Expanded human activity in, and development of, forested lands increase the interaction of both humans and domestic animals with wildlife. Among wild animals, black bears (*Ursus americanus*), which range across North America, have been significantly impacted by urban and suburban development (34). Such development has reduced forest habitat and increased forest fragmentation, leading to increased interactions between humans and bears, as evidenced by observations and property damage (35). Recently, the foodborne bacterial pathogen *Listeria monocytogenes* was found to be frequently isolated from black bears (36), and it is possible that bears may act as reservoirs and vehicles for other bacterial pathogens, including *C. jejuni*.

In the current study, we investigated the complete genome sequences of *C. jejuni* isolates collected from American black bears in the southeastern United States. Using these sequences, we conducted *in silico* multilocus sequence typing and performed comparative genomic analyses to determine similarity and phylogeny to worldwide *C. jejuni* isolates and those associated with the southeastern United States. Upon identifying two isolates with novel sequence types (STs), we explored their genetic content and identified genetic loci that are similar to loci in host-specific *C. jejuni* strains including *C. jejuni* subsp. *doylei*. These results identify American black bears as a new host and will help address the role that wild animals have in the epidemiology of *C. jejuni*.

## Materials and Methods

### Isolates and Culturing

Nine strains of *C. jejuni* were isolated from American black bear fecal and rectal swabs that had also been used to identify *Listeria monocytogenes*, all sampling locations, sample collection protocols, and permits have previously been described in detail in Parsons *et al* (36). Briefly, the bears lived in the southeastern United States, specifically Virginia, North Carolina, and Georgia, and the bear capture and/or handling protocols were approved by the Institutional Animal Care and Use Committee at North Carolina State University (14-019-O), the University of Georgia (A2011 10-004-A1) and Virginia Tech (12-112 and 15-162). Specifically for *Campylobacter* isolation, the samples (100 mg feces or the entire swab tip) were enriched for general *Campylobacter* spp. by resuspension in 10 mL Bolton broth (Oxoid Ltd., Hampshire, UK) and incubation at 37°C for 24 h under microaerobic conditions generated with GasPak EZ Campy sachets (Becton, Dickinson and Co., Sparks, MD, USA). The enrichments (100 μL) were plated for thermophilic *Campylobacter* on modified charcoal cefoperazone deoxycholate agar (mCCDA; Oxoid) with CCDA Selective Supplement (SR0155E, Oxoid), and the plates were incubated microaerobically at 42°C for 48 h, as described previously. Following incubation, an average of five putative *Campylobacter* colonies per sample were purified on Mueller-Hinton agar (MHA; Becton, Dickinson, and Co.), as described previously (37). *Campylobacter* confirmation was conducted via PCR using primers targeting *hipO*, as previously described (38).

### Bacterial isolate genome sequencing

DNA was extracted from *C. jejuni* isolates (**Table 1**) as described previously (39). Sequencing was carried out using the PacBio RS II (Pacific Biosciences, Menlo Park, CA) and Illumina MiSeq platforms (Illumina Inc., San Diego, CA). For the PacBio platform, SMRTbell libraries were prepared from 10 μg of bacterial genomic DNA with fragmentation in G-TUBEs (Covaris, Woburn, MA), following a BluePippin 10-kb size selection with a 0.75% DF Marker S1 high-pass 6- to 10-kb vs3 cassette (Sage Science, Beverly, MA) with 1× AMPure cleanup and DNA repair (40). Single-molecule real-time (SMRT) cells were run with 0.1 nM on-plate concentration, P6/C4 sequencing chemistry, the MagBead OneCellPerWell v1 collection protocol, and 360-min data collection mode. PacBio DNA internal control complex P6 was used as an internal sequencing control, and the read quality control was conducted using FastQC (Pacific Biosciences). The PacBio reads were assembled using the RS Hierarchical Genome Assembly Process (HGAP) v3.0 in SMRT Analysis v2.2.0 (Pacific Biosciences).

**Table 1.** Strain data, including MLST alleles, sequence types and clonal complexes from black bear *C. jejuni* isolates.

| Strain | Year | State | MLST alleles |  |  |  |  |  |  | ST | CC |
| --- | --- | --- | --- | --- | --- | --- | --- | --- | --- | --- | --- |
|  |  |  | *aspA* | *glnA* | *gltA* | *glyA* | *pgm* | *tkt* | *uncA* |  |  |
| SKBC1 | 2014 | North Carolina | 2 | 21 | 5 | 2 | 59 | 1 | 5 | 222 | 206 |
| SKBC3 | 2014 | North Carolina | 255 | 530 | 279 | 607 | 740 | 585 | 276 | 7630 |  |
| SKBC5 | 2014 | North Carolina | 305 | 756 | 279 | 607 | 479 | 585 | 276 | 10620 |  |
| SKBC9 | 2014 | Georgia | 12 | 6 | 61 | 147 | 261 | 32 | 3 | 10501 |  |
| SKBC25 | 2015 | North Carolina | 1 | 6 | 61 | 244 | 797 | 32 | 3 | 10624 | 179 |
| SKBC41 | 2015 | Virginia | 4 | 7 | 10 | 4 | 1 | 7 | 1 | 45 | 45 |
| SKBC57 | 2016 | North Carolina | 2 | 1 | 1 | 3 | 2 | 1 | 5 | 21 | 21 |
| SKBC94 | 2016 | North Carolina | 26 | 2 | 9 | 51 | 8 | 46 | 21 | 682 | 682 |
| SKBC102 | 2016 | North Carolina | 4 | 7 | 10 | 4 | 1 | 7 | 1 | 45 | 45 |

For the Illumina platform, libraries were prepared with the KAPA LTP library preparation kit (Roche) with Standard PCR Amplification Module (Roche), following the manufacturer’s instructions except for the following changes: 750 ng DNA was sheared at 30 psi for 40 sec and size selected to 700-770 bp following Illumina protocols. Standard desalting TruSeq LT primers were ordered from Integrated DNA Technologies (Coralville, IA) and used at 0.375 µM and 0.5 µM final concentrations, respectively. PCR amplification was reduced to 3-5 cycles. Libraries were quantified using the KAPA Library Quantification Kit, except with 10 µL volume and 90 sec annealing/extension PCR, then pooled and normalized to 4 nM. Pooled libraries were re-quantified by ddPCR on a QX200 system (Bio-Rad), using the Illumina TruSeq ddPCR Library Quantification Kit and following manufacturer’s protocols, except with an extended 2 min annealing/extension time. The libraries were sequenced on a MiSeq instrument (Illumina) using a MiSeq Reagent Kit v2 (500-cycles) at 13.5 pM, following the manufacturer’s protocols.

A final base call validation of the PacBio contigs was performed by mapping Illumina MiSeq reads that were trimmed to quality score threshold of 30 (Q30) or higher using the reference assembler within Geneious Prime 2020.2.1 (Biomatters, Ltd., Auckland, New Zealand). Single nucleotide polymorphisms (SNPs) between the PacBio assembly and the MiSeq reads were addressed using the annotate and predict/find SNPs module, with a minimum coverage parameter of 50 and a minimum variant frequency parameter of 0.8. The final corrected assemblies were annotated for protein-, rRNA-, and tRNA-coding genes using the NCBI Prokaryotic Genomes Annotation Pipeline (PGAP) (41). The chromosomal and plasmid sequences for each strain were deposited into NCBI under BioProject PRJNA971184.

### Determination of Penner capsular types, lipooligosaccharide locus classes, flagellar modification loci and *C. jejuni* integrated elements

Capsular polysaccharide (CPS)/Penner type, lipooligosaccharide (LOS) locus types, flagellar modification (FM) loci, and *C. jejuni* integrated elements (CJIEs) were identified by BLASTN analysis using Geneious Prime 2022.2. For CPS/Penner serotype determination, the *in silico* sequences of 36 PCR products for specific Penner types, as described previously (42, 43) were used in a BLASTN screen. For LOS locus classes, the variable sequences for LOS classes A-S (44, 45) were used in BLASTN searches of the nine genomes. For each *C. jejuni* genome, the LOS locus was compared to the LOS class exhibiting the closest match by BLASTN using Mauve. The genomes were screened for CJIEs, including the four elements identified within strain RM1221 (46) and the element containing the type VI secretion system (47, 48) using the BLASTN plugin in Geneious. For CJIEs and FM loci, the complete loci and individual genes were aligned using MAFFT and visualized using Mauve software to determine rearrangements, insertions, and deletions (49).

### Molecular typing and comparison

The nine complete genomes were submitted to the PubMLST database (https://pubmlst.org/organisms/campylobacter-jejunicoli) for curation and analysis. The seven-locus MLST sequence types (STs) were assigned as described previously (50, 51). Novel alleles and profiles for two strains, SKBC3 and SKBC5, were assigned new allele numbers and sequence types.

### Comparative Genomic Analyses

The black bear *C. jejuni* genomes were compared to other thermotolerant *Campylobacter* spp., including other *C. jejuni* genomes. To determine average nucleotide identity (ANI) values between *Campylobacter* spp., we used JSpecies (v. 1.2.1) with default parameters (52). The relationship between the *C. jejuni* isolates from black bears and a variety of *C. jejuni* isolates (**Table 2** and **Supplementary Table 1**) was explored using maximum likelihood phylogenies constructed from core genome alignments. Concatenated sequences of 1,107 core genes (present in 99% or more isolates) from 223 total *C. jejuni* genomes including from the following sources: bear isolates (n=9), cattle (n=48), poultry (n=90), environment (n=6), guinea pig specific (n=1), human clinical (n=35), monkey (n=2), sheep (n=1), swine (n=4), mouse (n=1), turkey (n=3), vole (n=1), water (n=15), wild bird (n=1), *C. jejuni* subsp. *doylei* (n=4) isolates, and unknown sources (n=2). Additionally, 143 of these genomes were from sources isolated in the southeastern United States particular Georgia, North Carolina, and Virginia to match where the bear isolates were also collected during the study. Core gene sequences were aligned using the MAFFT module (53) within Roary (v3.12.0), using the following flags: -e (create a multiFASTA alignment of core genes using PRANK); -n (fast core gene alignment with MAFFT); -v (verbose output to STDOUT); and -i 90 (minimum percentage identity for BLASTP; 90%)(54). Maximum likelihood phylogenies were constructed using Randomized Accelerated Maximum Likelihood (RAxML) with a GTRCAT model with 1,000 bootstraps (55) and visualized in Microreact (56). Additional phylogenetic comparisons were visualized using iTOL (51, 57).

**Table 2.** Genomic features of black bear *C. jejuni* isolates.

| Strain | Genome size (Mbp) | Genome acc. # | Penner | Variable genomic loci |  |  |  | Integrated genomic elements |  |  |  |
| --- | --- | --- | --- | --- | --- | --- | --- | --- | --- | --- | --- |
|  |  |  |  | LOS class | *cdtABC* | E-D | AMR<sup>c</sup> | CJIE1-Mu-like | CJIE2 | CJIE3 | CJIE4 |
| SKBC1 | 1.724 | CP125383 | unknown | C | Y | N | KTB | Y | Y | N | N |
| SKBC3 | 1.765 | CP125394 | HS:4cx | B2 | N | Y | - | Y2<sup>d</sup> | Y | N | N |
| SKBC5 | 1.784 | CP125391 | HS:16 | A2 | N | Y | - | Y2 | Y | N | N |
| SKBC9 | 1.676 | CP125390 | HS:3,17 | O | pseudo<sup>a</sup> | N | B | N | N | N | Y |
| SKBC25 | 1.708 | CP125388 | HS:1,44 | A2 | Y | N | B | Y | N | N | N |
| SKBC41 | 1.638 | CP125386 | HS:37 | H | Y | N | TB | N | N | N | N |
| SKBC57 | 1.681 | CP125396 | HS:2 | C | Y | N | B | N | N | N | Y |
| SKBC94 | 1.595 | CP125395 | HS:1,44 | Q | p/2<sup>b</sup> | N | B | N | N | N | N |
| SKBC102 | 1.651 | CP125387 | HS:55 | E | Y | N | TB | N | N | N | N |

- a. Pseudo: *cdtABC* genes present with point mutations. b. p/2, *cdtABC* genes present with point mutations and a second *cdtABC* operon is present. c. K, presence of *aph3*; T, presence of *tet(O)*; B, *blaOXA*; -, no AMR markers. d. Y2, two Mu-like prophages present

Gene families were identified using Roary software at 90% identity across the entire gene, which generated core (in <99% - 100% of all the genomes), soft-core (in <95% - 99% of all the genomes), shell (<15% - 95% of all the genomes), and cloud (>0% - 15%) genes, and visualized using the roary_plots.py script.

### Phylogenetic dendrogram construction of Entner-Doudoroff loci

The Entner-Doudoroff (E-D) locus from *C. jejuni* strain SKBC3 (CP125394 from nucleotide 39,023-47,678) was used as bait to identify E-D loci in other *Campylobacter* genomes using BLASTN specific to the *Campylobacter* genus within the NCBI non-redundant (nr) database. An E-D locus was previously identified in a *C. jejuni* strain (GP012) isolated from guinea pigs; however, this locus is not in the NCBI nr database and was added separately. Loci were obtained from the following strains: CP000768 (strain 269.97), CP059375 (2010D-8469), CP125391 (SKBC5), CP125394 (SKBC3), JACRSF000000000 (GP012) and LR134359 (NCTC11951^T^) from *C. jejuni and C. jejuni* subsp. *doylei*; CP017875 (ZV1224) and KT001110 (CHW475) from *C. coli*; CP031611 (NCTC 13823^T^), CP063536 (USA52) and CP065357 (UF2019SK1) from *C. hepaticus*; CP059597 (LMG 32306^T^) from *C. molothri* (58); and CP020867 (LMG 24588^T^) from *C. cuniculorum* were aligned using MAFFT, and a dendrogram was constructed within MEGA v.11 using the Maximum Likelihood method and Tamura-Nei model (59–61).

## Results

### *C. jejuni* isolates from American black bears are genotypically diverse

We sequenced the genomes of nine strains of *Campylobacter jejuni* isolated from 2014 to 2016 from American black bears living in the southeastern United States (**Table 1**). Eight sequence types (ST) were assigned to these nine *C. jejuni* genomes by PubMLST (**Table 1**). Six of the nine strains were members of previously identified *C. jejuni* clonal complexes. These included CC 21, CC45, CC206, CC179 and CC682, as identified in the strains SKBC57, SKBC41 and SKBC102, SKBC1, SKBC25 and SKBC94, respectively. *C. jejuni* within clonal complexes CC21, CC45 and CC206 have been described previously as host generalists (13). Similarly, the non-human isolation sources of *C. jejuni* within CC-179 includes poultry, cattle and environmental water (Supp. Table 2). It should be noted that strain SKBC9 assigned to ST-10501 and no clonal complex, shares four alleles with ST-179 complex isolate SKBC25. These alleles were shared by 70 strains in PubMLST (**Supplementary Table 2**). The strain SKBC94 (ST-682 complex) would be considered a species-specific *C. jejuni* with 165/205 ST-682 complex strains from PubMLST isolated from European starlings (*Sturnus vulgaris*) (**Supplementary Table 3**).

Strains SKBC3 and SKBC5 were assigned novel sequence types, ST-7630 and ST-10620, respectively. These two strains share four alleles (*gltA279*, *glyA607*, *tkt585* and *uncA276*), and these alleles are represented at most by only one other strain in the database in Feb 2025 (**Supplementary Table 4**). Moreover, most isolates with shared alleles were from strains that were isolated from environmental waters in North America, except for the *aspA* allele that were also shared with two human isolates, a dog isolate, and an isolate from an unknown source (**Supplementary Table 4**).

### Genomic features of *C. jejuni* isolated from black bears

The genome sizes of these *C. jejuni* bear isolates were quite variable with estimated chromosome sizes ranging from 1.595-1.784 Mbp (**Table 2**) and containing between 1,491 and 1,689 predicted protein-encoding genes. The pangenome for these nine *C. jejuni* chromosomes was 3,356 genes with 1,256 core genes and 2,100 accessory genes (present in >90% of genomes). Three strains (SKBC1, SKBC5 and SKBC25) have plasmids that were not included in the pangenome analysis. Major plasmid features are described below.

The differences in chromosome sizes between the strains was mostly the result of the presence or absence of *Campylobacter jejuni* integrated elements (CJIEs). CJIEs, which include bacteriophages, plasmid-like elements and transposons have been identified in many *C. jejuni* strains (46–48, 62–64). The strains with the largest genomes, SKBC1, SKBC3 and SKBC5 (1.724, 1.765 and 1.784 Mb, respectively), possess multiple CJIEs. Bacteriophages similar to the Mu-like phage CJIE1, CJIE2, and CJIE4 were each detected by BLASTN in at least one bear isolate (**Table 2**).

Mu-like prophage (CJIE1-like) are harbored by four strains (SKBC1, SKBC3, SKBC5 and SKBC25) (**Table 2**, **Supplementary Figure 1** and **Supplementary Table 5**). These Mu-like prophages are integrated in different genomic locations in these strains (**Supplementary Table 5**) and were classified into two types based on the presence or absence of *dns*, a gene encoding an extracellular DNase, and major differences in sequence identity of genes encoding DNA transposition protein A, DNA transposition protein B, and Mu-like bacteriophage tail structural and assembly proteins (**Supplementary Figure 1** and **Supplementary Table 5**). Strains SKBC3 and SKBC5 each harbor both types of Mu-like bacteriophages, whereas SKBC1 and SKB25 only harbor one of the two Mu-like bacteriophages. Strains SKBC3, SKBC5 and SKBC25 possess Mu-like bacteriophages containing *dns*. These bacteriophages have >82% identity to CJIE1 from *C. jejuni* strain RM1221 (**Supplementary Figure 1A**). The other three Mu-like bacteriophages, in strain SKBC1 (near *npdA*) and the second Mu-like bacteriophages in strains SKBC3 (near *ctsT*) and SKBC5 (tRNA-Arg gene near *aroB* and *tgt*) do not harbor *dns,* and are similar only for certain viral genes within RM1221 CJIE1, specifically, genes encoding the host-nuclease inhibitor (Gam), the viral baseplate and some viral tail-related proteins (**Supplementary Figure 1B** and **Supplementary Table 5**).

The other two bacteriophage families, CJIE2-like and CJIE4-like, are present in more than one of the isolates from the black bears. These bacteriophage families have specific genomic integration sites, with CJIE2-like bacteriophages inserted at the tRNA-Arg gene adjacent to *fusA* and CJIE4-like bacteriophages inserted at the tRNA-Met gene adjacent to *rodA.* The CJIE2-like bacteriophages are present in strains SKBC1, SKBC3 and SKBC5. Comparison of these CJIE2-like bacteriophages with CJIE2 in strain RM1221 revealed diversity, identifying large insertions and deletions of genes (**Supplementary Figure 2**). The CJIE4-like bacteriophages are present in strains SKBC9 and SKBC57 and are >98% identical to CJIE4 from strain RM1221 (**Supplementary Figure 3**).

Other insertion elements were also observed in the *C. jejuni* from black bears. Although there is an integrated element at the CJIE3 integration site (tRNA-Arg gene near *aroB* and *tgt*) in strain SKBC5, this element is quite distinct from the RM1221 CJIE3 with only similarities at the site-specific recombinase (integrase) and TraG-like genes that may have a role in integration site selection and conjugation, respectively (**Supplementary Figure 4**). A CJIE, containing a type VI secretion system (T6SS), possessed by some *C. jejuni* (47, 48) is not present as an integrated element in any of the nine bear isolates; however, strain SKBC25 possesses a plasmid (pSKBC25) with T6SS genes.

Besides differences in genome sizes due to CJIEs, the large hypervariable genomic loci encoding major surface structures, including capsule biosynthesis, lipooligosaccharide biosynthesis and flagellar modification (42, 44, 65), are mostly distinct between the nine strains. As shown in Table 2, there are eight different capsular biosynthesis loci (Penner serotypes) and seven different LOS biosynthesis loci. Three LOS loci (A2, B2 and C) in five strains are potentially capable of synthesizing sialylated LOSs that have been associated with post infection neuropathies (66). All nine FM regions are distinct and ranged in size from 27kb to 50kb (**Supplementary Figure 5**).

Other notable variable regions included transposable elements, plasmids and chromosomal loci defined as hypervariable by comparative genomic indexing (64, 65). Strains SKBC41 and SKBC102 each possesses an IS607-like transposable element, containing the tetracycline resistance gene *tet(O)*, inserted near *rarA*. Only three strains possess plasmids, and none are shared between strains. Strain SKBC1 carries two plasmids: pSKBC1-1 harboring *tet(O)* and *aph(3’)-IIIa* and pSKBC1-2. Both pSKBC1-1 and pSKBC1-2 show >98% identity to multiple plasmids. Strain SKBC5 carries two plasmids: pSKBC5-1 harboring putative bacteriocins and pSKBC5-2. pSKBC5-1 is fairly novel, with only a few plasmids identified by BLASTN that aligned with portions of pSKBC5-1. pSKBC5-2 shows >98% identity to pTet-like plasmids but does not harbor the *tet(O)* gene. Finally, SKBC25 carries pSKBC25, which harbors a T6SS, as mentioned above.

Variation among the bear strains includes regions that are variably present on the chromosome including pantothenate (vitamin B5) biosynthesis locus (*panBCD*), *ggt* that encodes gamma-glutamyltranspeptidase, which plays a role in metabolism of glutathione and glutamine, and *metAB* (*metA*X) that encode enzymes involved in converting L-homoserine to L-methionine (64, 67, 68). The *panBCD* and *metAB* loci were detected in the same two strains (SKBC1 and SKBC57), while *ggt* was detected in four strains (SKBC3, SKBC5, SKBC41 and SKBC102), none of which overlapped with possession of *panBCD* and *metAB*. Antibiotic resistance genes were present in some of the bear isolates (Table 2). Strains SKBC41 and SKBC102 possess *tet(O)* on the chromosome and strain SKBC1 harbors *tet(O)* and *aph(3’)-IIIa* on a plasmid, as mentioned above. It should be noted that each of these strains belong to MLST clonal complexes that are associated with domestic animals. Seven of the isolates, SKBC1, SKBC9, SKBC25, SKBC41, SKBC57 and SKBC102, possess a *bla*_OXA_ gene near *modC*. None of the strains have mutations in *gyrA* or the 23S RNA gene that confer resistances to fluoroquinolones and macrolides, respectively. Strains SKBC3 and SKBC5 have no identifiable antimicrobial resistance markers and no evidence of a remnant of a *bla*_OXA_ gene near *modC*.

### ANI and phylogenetic analysis also support carriage of both common and distinct *C. jejuni* strains

The genomic relationship of the black bear isolates with several other thermotolerant *Campylobacter* was examined using average nucleotide identity (ANI) analysis. Within the collection here, all *C. jejuni* strains have ANI values >94% against all other *C. jejuni* strains, while the ANI values are <85% against *C. coli* and *C. hepaticus*, and <76% with *C. upsaliensis*, *C. vulpis* and *C. helveticus* (**Figure 1**). Among the bear isolates, seven strains (SKBC1, SKBC9, SKBC25, SKBC41, SKBC57, SKBC94, SKBC102) have ANI values between 96.9% and 99.1% against all other *C. jejuni* subsp. *Jejuni* strains. However, the bear isolates SKBC3 and SKBC5 have ANI values <97% and <96% against the other *C. jejuni* subsp. *jejuni* and *C. jejuni* subsp. *doylei* strains, respectively. The ANI values between the two subspecies, *C. jejuni* subsp. *jejuni* and *C. jejuni* subsp. *doylei*, were between 95 and 96% (**Figure 1**). Thus, SKBC3 and SKBC5 were slightly distinct from other *C. jejuni* subsp. *jejuni* but not as much as *C. jejuni* subsp. *doylei* strains, or the guinea pig-associated *C. jejuni* strain that had ANI values <95% to all other *C. jejuni*.

**Figure 1.**
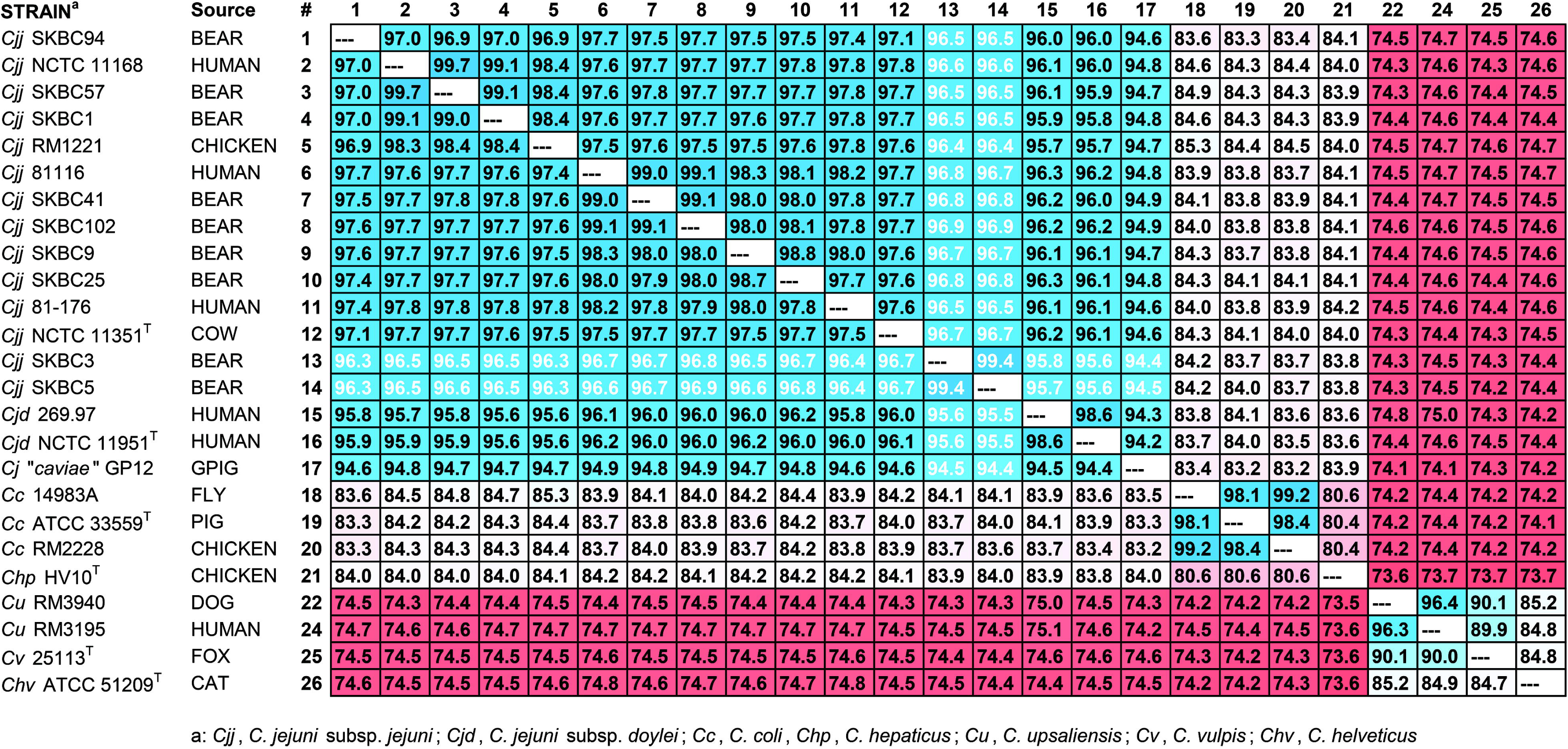
Average Nucleotide Identity (ANI) Values. Note—Values represent averages (in %) of each pair. Values ≥ 90% are shaded in blue; values between 81 and 89% are shaded in white grey and values < 81% are shaded in red.

Pan-genome analyses were conducted using Roary analysis on the nine bear and with 71 global *C. jejuni* genomes, including *C. jejuni* subsp. *jejuni,* a strain isolated from a guinea pig, a bank vole, isolates from water, and *C. jejuni* subsp. *doylei* and an additional 143 *C. jejuni* genomes isolated from various sources (e.g. poultry, environment waters, cattle, etc.) in the same southeastern United States region as the bear isolates were obtained (**Supplementary Table 1**). All genomes were downloaded from NCBI and used to gain a better appreciation of the genomic differences of the strains at the individual gene level. Roary analysis was performed at a minimum percentage identity for BLASTP of 90% since the ANI values for several strains were below 95%. At the 90% cutoff, there were 1,107 core genes found in >99%, 236 soft core genes in =>95% to < 99%, 613 shell genes in =>15% <= and < 95% of strains and 3,328 cloud genes >0% to < 15% of strains.

Phylogenetic analysis compared the nine *C. jejuni* isolates from black bears with 71 global *C. jejuni* genomes, and 143 *C. jejuni* genomes from non-clinical strains isolated from the southeastern United States (Georgia, North Carolina and Virginia) (**Supplementary Table 1**). The analysis was performed using 1,107 *C. jejuni* core genes as determined by Roary analysis above. The resulting phylogeny demonstrated that these nine *C. jejuni* genomes from black bears were widely distributed among the other *C. jejuni* genomes (Figure 2). Bear isolates SKBC3, SKBC5, and SKBC94 were located on a major branch that includes *C. jejuni* subsp. *doylei* strains and the strains from a guinea pig, bank vole and certain environmental water samples (**Figure 2**). The other black bear *C. jejuni* isolates (SKBC1, SKBC9, SKBC25, SKBC41, SKBC57 and SKBC102) were in clusters comprised of other *C. jejuni* subsp. *jejuni* isolated from several animal sources (chicken, cattle, pigs, and turkey) and river water.

**Figure 2.**
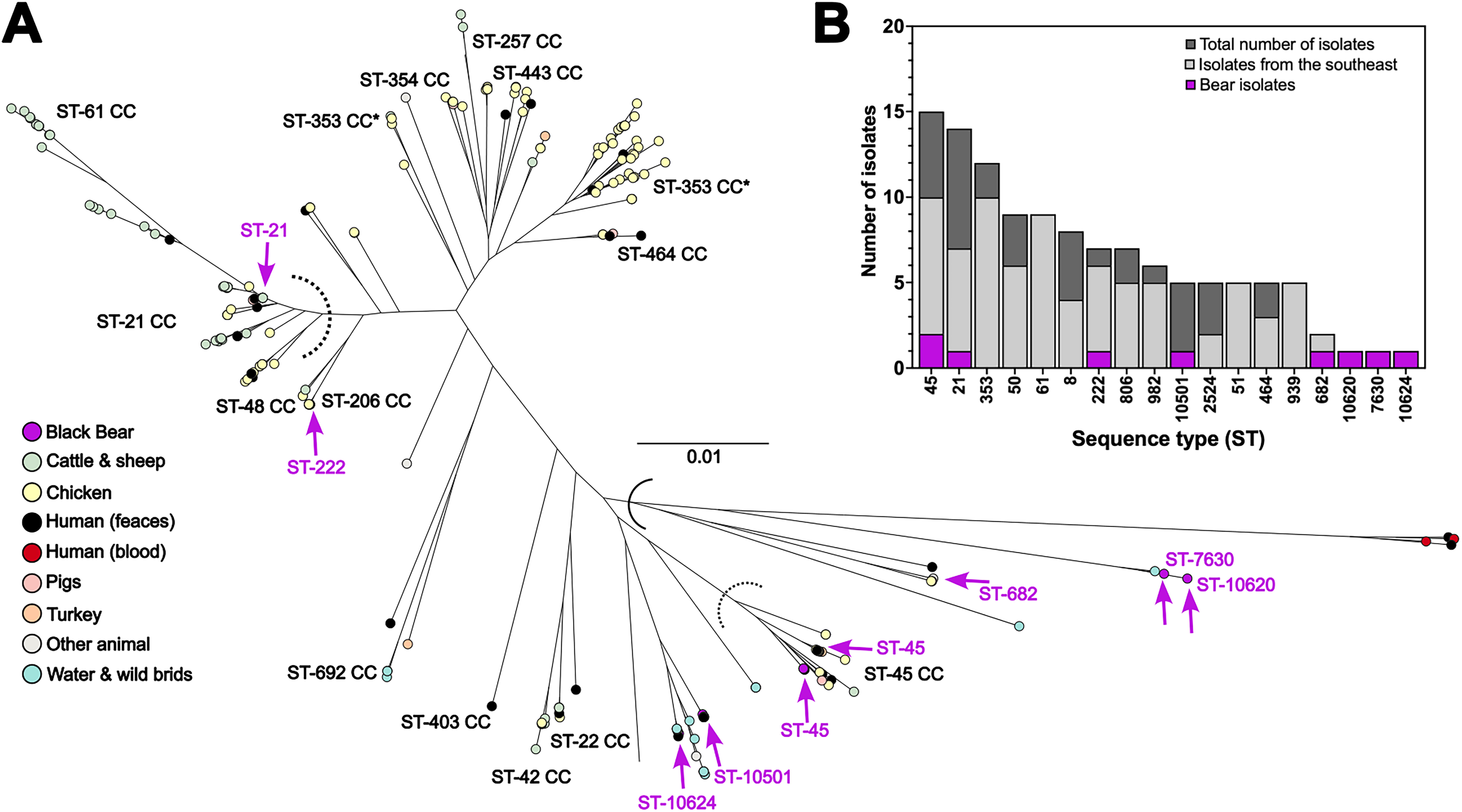
Phylogenetic and MLST genotypic distribution of *C. jejuni* isolates from black bears compared to a representative collection of *C. jejuni.* **A:** The relationship between isolates from black bears and a variety of *C. jejuni* isolates (**Table 1 and Supplementary Table 1**) was explored using a maximum likelihood phylogeny. Concatenated sequences of 1,107 core genes (present in 90% or more isolates) from 214 *C. jejuni* and *C. jejuni* subsp. *doylei* isolates were used to compare the nine bear isolates (purple nodes and indicated with arrows). Isolates from other diverse sources are colored by source, including black bear (n=9; blue), chicken and poultry farm (n=96; yellow), cattle and sheep (n=49; green), turkey (n=3; orange), water and wild birds (n=16; light blue), human feces (n=35; black), human blood (n=4; red), pigs (n=4; pink) and other animal sources (n=7; grey). Core gene sequences were aligned using the MAFFT module (53) within Roary (v3.12.0)(54). Maximum likelihood phylogenies were constructed using Randomized Accelerated Maximum Likelihood (RAxML), with a GTRCAT model with 1,000 bootstraps (55) and *C. jejuni* only topology is visualized in microreact (56) (https://microreact.org/project/tYPSuzXxiXyoUxa3R8ytvK-parker-et-alcampylobacter-from-us-black-bears) Major lineages are labeled (ST-clonal complexes), with dotted lines indicating the threshold between CCs **B:** MLST distribution of black bear isolates (purple shading) in this comparison dataset (dark grey) and those collected from the Southeast USA only (light grey) among sequence types (ST).

### Absence of the *cdtABC* and *mfrABE* loci within a phylogenetic cluster

In most *C. jejuni* subsp. *jejuni* strains, the *cdtABC* locus is adjacent to *lctP.* Strains SKBC1, SKBC25, SKBC41, SKBC57 and SKBC102 had a *cdtABC* locus predicted to be functionally complete at this location. Strains SKBC9 and SKBC94 had nonfunctional *cdtABC* loci adjacent to *lctP,* with both possessing a nonsense mutation in each *cdt* gene. However, in strain SKBC94, a second *cdtABC* locus was identified near the *lpxB* gene. This *lpxB-*linked *cdtABC* locus was observed in four of the other 214 *C. jejuni* genomes used in this study. The *lpxB-*linked *cdtABC* is the primary location of the *cdtABC* locus within *C. coli*, *C. lari* and other Campylobacters. BLASTN analysis of the *lpxB-*linked *cdtABC* locus from strain SKBC94 shows the closest alignments (∼77%) to *C. lari* strains. Strains SKBC3 and SKBC5 had major deletions within the *cdtABC* locus adjacent to *lctP*, with only remnants of *cdtC* remaining, and neither strain had the *lpxB-*linked *cdtABC*. It should be noted that all *C. jejuni* sharing the phylogenetic branch with bear isolates SKBC3, SKBC5, and SKBC94 had nonfunctional *lctP-*linked *cdtABC* loci (Supplementary Figure 6: orange and blue branches) Among these strains, those within the blue branch have major deletions of the *lctP-*linked *cdtABC* locus.

The *mfrABE* operon (previously *sdhABC*) that encodes methylmenaquinol:fumarate reductase was also mutated in this same cluster of “host specific” strains that includes strains SKBC3 and SKBC5, three *C. jejuni* subsp. *doylei* strains, and isolates from a guinea pig, bank vole, and water (**Supplementary Figure 6**: blue branch). Methylmenaquinol:fumarate reductase affects resistance to H_2_O_2_ (69). In SKBC3 and SKBC5, *mfrA* has a nonsense mutation and *mfrB* and *mfrE* are deleted. A variety of *mfrABE* deletions are found in other *C. jejuni* in this cluster. There are *C. jejuni* strains outside of this cluster containing mutated *mfrABE*; however, in these *C. jejuni*, the *mfrABE* mutations are nonsense mutations that could more easily revert to a functional methylmenaquinol:fumarate reductase system.

### Acquisition of the Entner-Doudoroff Pathway

The genomes of strains SKBC3 and SKBC5 possessed genes for the Entner-Doudoroff (E-D) pathway, which catalyzes the conversion of glucose-6-phosphate to pyruvate. These genes are situated within an rRNA locus between the 23S RNA gene and the 16S RNA gene. However, despite the E-D loci from SKBC3 and SKBC5 being 99.8% identical, the specific rRNA loci is distinct between SKBC3 and SKBC5. The E-D locus in SKBC3 is in the rRNA locus linked to *pyrG*-*recJ* while the E_D locus in SKBC is in the rRNA locus linked to *cfrA-hcrA*. Other *Campylobacter* possess E-D genes, and we performed alignment and maximum likelihood phylogenies to compare the E-D genes from the two bear-associated strains to E-D genes from *C. jejuni* isolated from a guinea pig, *C. jejuni* subsp. *doylei* and other *Campylobacter* species. Our tree demonstrates that the E-D loci from SKBC3 and SKBC5 are situated phylogenetically between E-D loci from *C. jejuni* and *C. coli.* (**Figure 3**).

**Figure 3.**
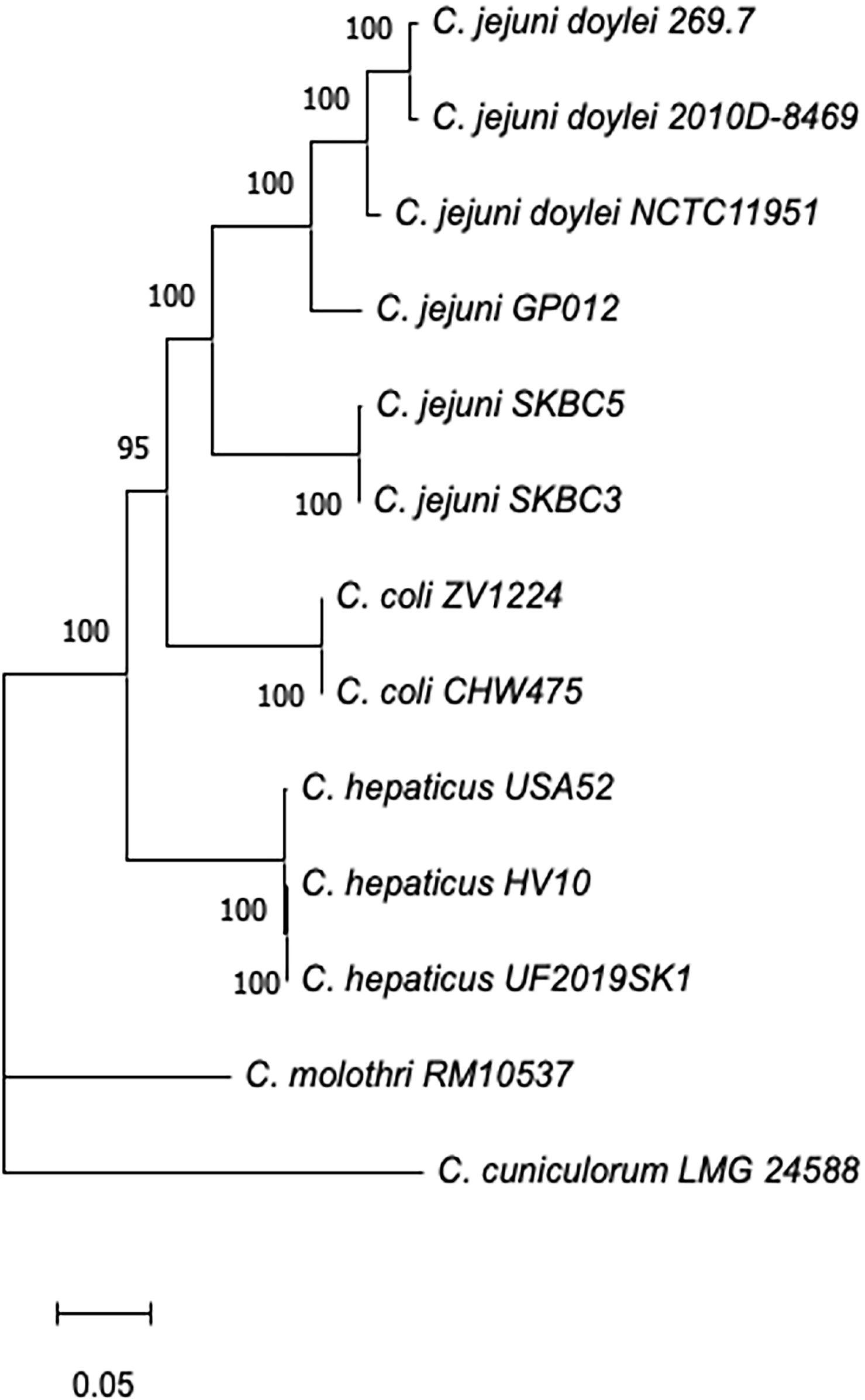
Phylogenetic characterization of the Entner-Doudoroff pathway locus. The phylogenetic tree of the E-D pathway locus from 13 *Campylobacter* strains. The dendrogram is drawn to scale, with branch lengths measured in the number of substitutions per site.

## Discussion

We isolated the foodborne pathogen *C. jejuni* from American black bears in the southeastern United States. Despite isolating only nine *C. jejuni* strains, these strains were from three different states and over a three-year period, suggesting *C. jejuni* commonly infects black bears. The nine *C. jejuni* strains were genotypically variable with eight distinct STs and six CCs. This *C. jejuni* strain variability may suggest that bears acquire the bacteria from both anthropogenic sources (trash receptacles or agricultural sites) and from non-human environments (water or wild animals).

Genomic comparisons of the isolates revealed substantial variability in chromosome sizes and gene content. The genome sizes ranged from 1.595 to 1.784 Mb and the variation was largely due to the presence of CJIEs as described previously (46–48, 62–64). For instance, strains SKBC1, SKBC3, and SKBC5 had the largest genomes due to multiple CJIEs (Table 2). The presence of these elements not only affects genome size but may also shape the functional potential of the strains by altering gene expression and natural competence (70–72). Differences in hypervariable loci encoding biosynthesis of surface glycan structures further emphasizes the genomic variability of these strains, with eight different CPS loci, seven different LOS loci and nine different FM loci identified in the nine strains. *Campylobacter* surface glycans, particularly CPS and LOS, are quite immunogenic and variable. The genetic variation at these glycan loci suggests that the bear isolates exhibit a variety of glycan surface structures. These glycans often play significant roles in virulence, with the best understood of these being the role of molecular mimicry of human gangliosides by particular sialylated LOS structures in the development of post-infection neuropathies (73). In fact, five of the *C. jejuni* strains from bears possessed LOS loci that could potentially synthesize sialylated LOS structures. Other variable loci include *panBCD*, *metAB* (*metAX*), and *ggt*. These genes have been identified as playing a role in colonization of domestic cattle or poultry under certain conditions (67, 74–76); however, their variable presence in cattle and poultry isolates, as described previously (64, 65), and here, within bear isolates, suggests that they are not essential for the infection process in these hosts.

*C. jejuni* genomes are prone to recombination, and signals of host adaptation can be identified providing information of the strain’s recent ancestry (21, 77, 78). Based on their MLST CC, four *C. jejuni* strains isolated from the bears (SKBC1, SKBC41, SKBC57 and SKBC102) are predicted to be host generalists (21), according to their membership in the clonal complexes CC-21, CC-45, and CC-206. Host generalist *C. jejuni* are associated with broad host ranges, including domestic poultry and domestic cattle, and they are commonly recovered from human clinical samples (12, 79). Additionally, strains SKBC9 and SKBC25 are part of a group of over 70 strains that share four MLST alleles (Supplementary Table 2). The strains in this group are also likely host generalist, with nonclinical strains isolated from both domestic poultry and cattle, and environmental waters. In fact, these generalist strains from bears clustered with poultry and cattle strains within our phylogenetic analysis (Figure 2 and Supplementary Figure 6). Furthermore, three of these generalist strains possess antimicrobial resistance (AMR) genes. Surveys of avian *C. jejuni* strains suggested that *C. jejuni* from birds more closely associated with anthropogenic settings have a higher prevalence of AMR genes (80). Given the increased human activity and reduction of wild habitat where black bears live, there has been an increase in interactions between humans and bears (34, 35). Black bears generally stay away from humans; however, in agricultural areas where food is common, bears often feed on crops and can encounter domestic animals and their waste. Considering this, it is likely that black bears were colonized by generalist *C. jejuni* while foraging around agricultural areas or human habitations.

Three *C. jejuni* isolates from the black bear exhibited evidence that they were from non-agricultural environments. SKBC94 is in the ST-682 complex that is comprised mostly of strains from European starlings (*Sturnus vulgaris*). SKBC3 and SKBC5 currently have novel STs but share specific alleles with *C. jejuni* isolated from environmental waters, and thus unknown animal sources. Infection with *C. jejuni* from wild birds would not be unusual considering the observations of black bears eating wild bird nestlings (81). Black bears are also scavengers and could have obtained *C. jejuni* from a variety of dead animals or from environmental waters. Alternatively, strains SKBC3 and SKBC5 may represent bear-specific *C. jejuni*; however, there is not enough evidence to determine this currently.

All three of these strains (SKBC94 (ST-682), SKBC3 (ST-7630) and SKBC5 (ST-10620)) cluster phylogenetically with *C. jejuni* isolated from wildlife and environmental waters, and *C. jejuni* subsp. *doylei* strains (Figure 2 and Supplementary Figure 6). Beyond the phylogenetic clustering, these three bear isolates and the strains within the cluster (Figure 2 and Supplementary Figure 6) also had mutations within the *lctP-* linked *cdtABC* locus that would eliminate function. For SKBC94, each *lctP-*linked *cdt* gene had a nonsense mutation. On the other hand, SKBC3 and SKBC5 possessed major deletions of the *lctP-*linked *cdtABC* locus previously reported for *C. jejuni* subsp. *doylei* strains (82) and among *C. jejuni* subsp. *jejuni* strains from guinea pigs and wild birds (11, 83). Strains SKBC94 also possessed a second *cdtABC* operon linked to *lpxB* that was shared by multiple CC ST-682 strains. The absence of *lctP-*linked *cdtABC* and/or the presence of *lpxB-*linked *cdtABC* suggests that cytolethal distending toxin may play a major role in host-specific colonization as noted by Guirado (83). Additionally, the absence of *mfrABE* in SKBC3, SKBC5 and other strains in the cluster suggests methylmenaquinol:fumarate reductase may play a negative role in host-specific colonization. Methylmenaquinol:fumarate reductase may contribute to the generation of oxidative stress in *C. jejuni* by creating reactive oxygen species. *C. jejuni* mutants unable to produce methylmenaquinol:fumarate reductase were demonstrated to be more resistant to H_2_O_2_ (69). It is not exactly clear how either cytolethal distending toxin or methylmenaquinol:fumarate reductase affects host specificity. More studies will be required to elucidate their roles.

Finally, we identified the presence of the E-D locus in strains SKBC3 and SKBC5. The E-D locus encodes enzymes involved in the Entner-Doudoroff pathway that catalyze the conversion of glucose to pyruvate and has been identified in *C. jejuni*, *C. coli* and other *Campylobacter* species (11, 84). Vegge et al (84) demonstrated the fitness advantages within *C. coli* possessing the E-D locus including stationary-phase survival and biofilm production. Despite the physiological advantages, the E-D locus was in less than 2% of *C. jejuni*/*C. coli* strains. Moreover, we found that the E-D loci from the bear-associated strains are distinct from E-D from other *C. jejuni*. The diversity of E-D loci was previously observed by Vegge et al for 113 *Campylobacter* isolates and they suggested this may be the result of different times of E-D introduction into each strain followed by evolution (84). Alignment and maximum likelihood trees demonstrate that the E-D loci in SKBC3 and SKBC5 are situated phylogenetically between E-D loci from *C. jejuni* (e.g. E-D loci in a guinea pig-associated *C. jejuni* strain and loci from three of *C. jejuni* subsp. *doylei* strains) and E-D loci from *C. coli*.

## Conclusions

This study demonstrates for the first time that *C. jejuni* can be isolated from American black bears. This is not the only human pathogen that has been isolated from bears. *Listeria monocytogenes* was also identified in feces, rectal and nasal swabs of black bears (36). Due to the bears attraction to food sources near agricultural settings, we suggest that black bears could serve as a natural vehicle and perhaps a reservoir for a wide variety of *C. jejuni* lineages from both domestic animals and wildlife. The limited number of *C. jejuni* strains does not permit assessments of the overall prevalence of *C. jejuni* carriage in black bears or allow rigorous genome-wide association studies.

However, we distinguished both domestic animal-associated genotypes and wild bird genotypes. Moreover, we identified two strains with novel genotypes that were only associated with black bears. Thus, sequencing and phenotypic analysis of additional *C. jejuni* isolates from black bears will be necessary to determine if these such strains are bear specialists or specialists for another host.

## Contributors

Conceptualization: CTP, SK and KKC. SK led collection of the isolates. SH sequenced the isolates. Formal Analysis: BP, KKC, CTP, WGM, SH. Sample collection: JN, CD, NG. Original Draft Preparation: CTP, SK, KKC. All authors read and approved the final manuscript.

## Acknowledgements

This research was also supported in part by USDA-ARS CRIS project 2030-42000-055-00D. This research was partially funded by the USDA National Institute of Food and Agriculture, award 2018-67017-27927, and Technology and Research Initiative Fund (TRIF) provided to Kerry Cooper by the University of Arizona. No funding agency had any role in the study design, data collection and analysis, decision to publish, or preparation of the manuscript. We wish to thank Jeff Niedermeyer for culturing the *C. jejuni* strains from the bear samples.

## Abbreviations

AMR: antimicrobial resistance
CC: clonal complex
MLST: multi-locus sequence typing
ST: sequence type
WGS: whole genome sequencing

