## Supplementary Figures for "American black bear (*Ursus americanus*) as a potential host for *Campylobacter jejuni*"

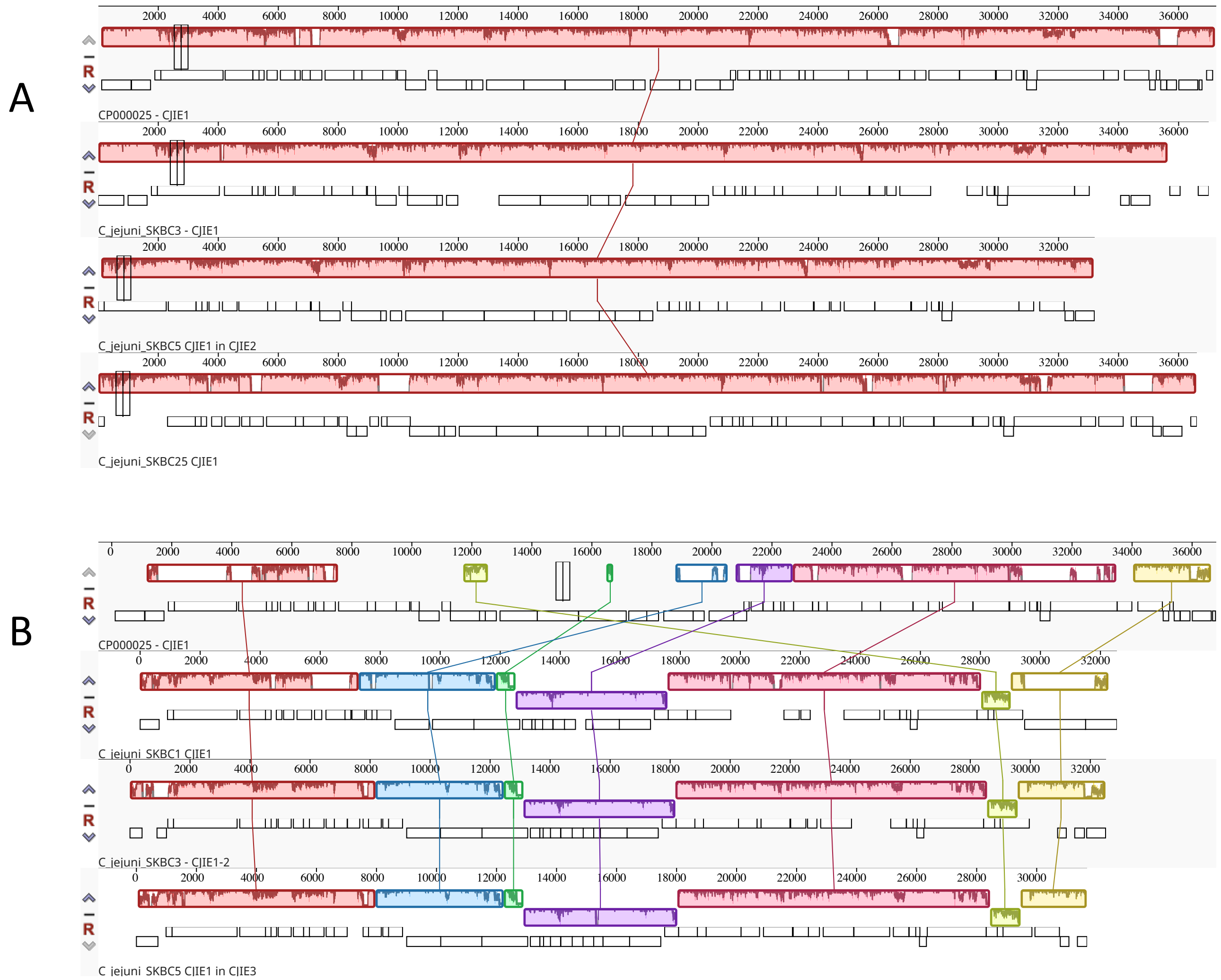


**Supplementary Figure 1. Genome alignment of CJIE1 (Mu-like bacteriophages).** Genome alignment of CJIE1-like genomes from Campylobacter jejuni strain RM1221 and the bear isolates with CJIE1-like bacteriophages with similar transposase-encoding genes (A) using Mauve revealed one collinear block conserved among bacteriophage genomes disrupted by insertions and deletions. Genome alignment of CJIE1-like genomes from Campylobacter jejuni strain RM1221 and the bear isolates with CJIE1-like bacteriophages with dissimilar transposase-encoding genes (B) revealed seven collinear blocks conserved among bacteriophage genomes disrupted by insertions and deletions. Each bacteriophage genome is arranged horizontally and homologous blocks in each genome are shown as identically colored regions linked across the Mu-like bacteriophages. Conserved blocks that were inverted compared to RM1221 in the figure are located beneath the CJIE1-like bacteriophage genome. The order of Mu-like bacteriophages with similar transposase-encoding genes to RM1221 (A) is: RM1221, SKBC3, SKBC5 and SKBC25. The order of Mu-like bacteriophages with dissimilar transposase-encoding genes to RM1221 (B) is: RM1221, SKBC1, SKBC3, and SKBC5.

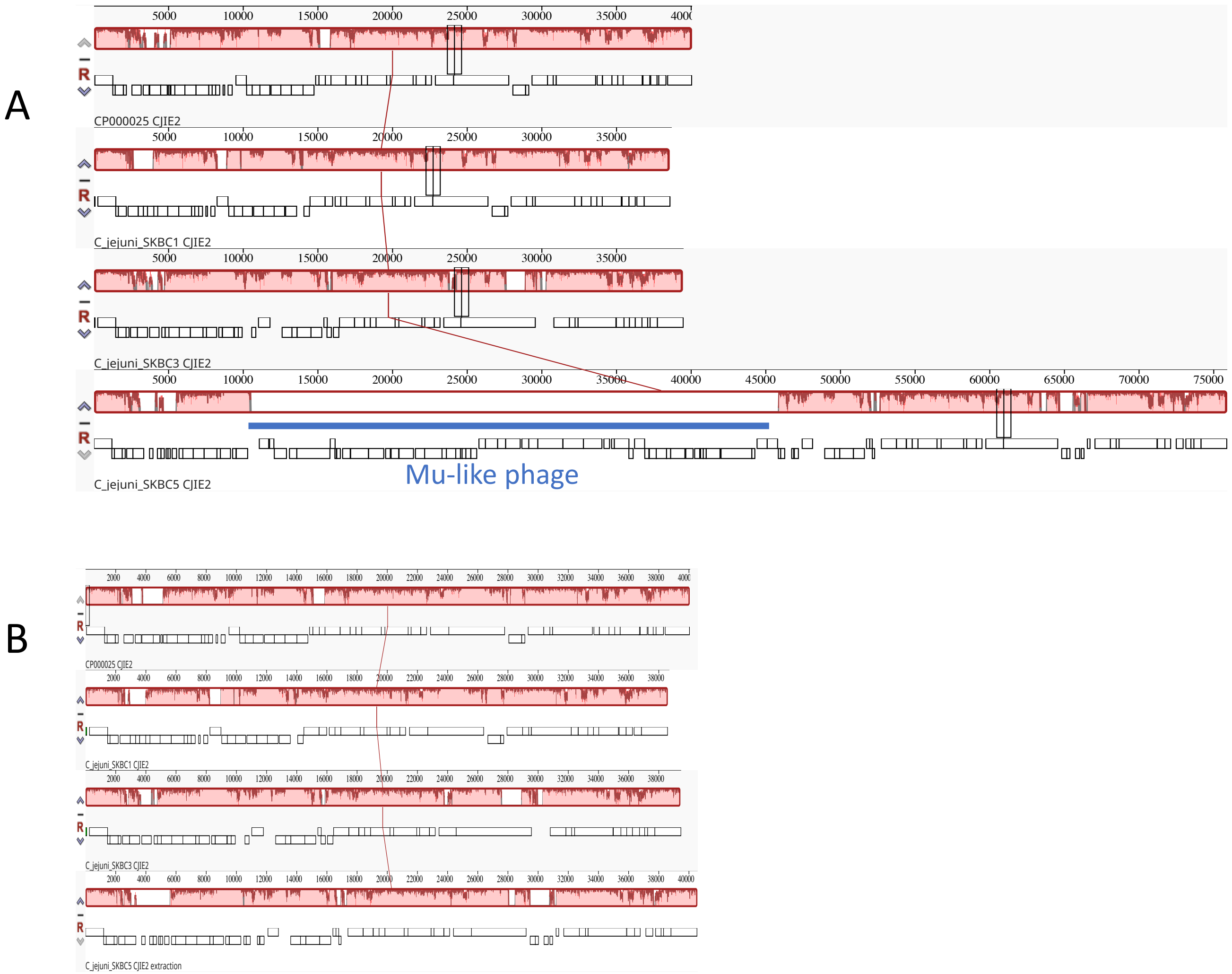


**Supplementary Figure 2. Genome alignment of CJIE2.** Genome alignment of CJIE2-like genomes from C. jejuni strain RM1221 and the C. jejuni isolates from bears using Mauve revealed one collinear block conserved among bacteriophage genomes disrupted by insertions and deletions (A). The order of CJIE2-like genomes is RM1221, SKBC1, SKBC3, and SKBC5. CJIE2 in SKBC5 possesses a Mu-like bacteriophage is underlined with a blue bar. This Mu-like bacteriophage element disrupts the collinearity of the CJIE2-like genomes. In (B), genome alignment of CJIE2-like genomes from C. jejuni strain RM1221 and the C. jejuni isolates from bears using Mauve, in which the Mu-like bacteriophage has been removed from the CJIE2-like bacteriophage of SKBC5.

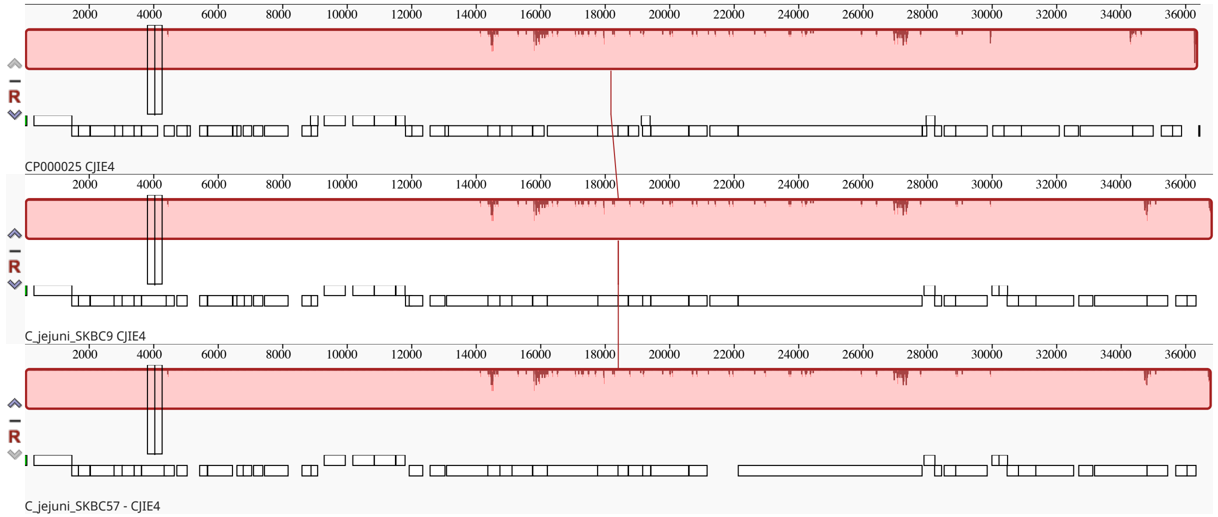


**Supplementary Figure 3. Comparisons of CJIE4.** Genome alignment of CJIE4 genomes from *C. jejuni* strain RM1221 and the *C. jejuni* isolates from bears using Mauve revealed one collinear block conserved among bacteriophage genomes with no large insertion or deletion disruptions. The order of aligned CJIE4 is: RM1221, SKBC9, and SKBC57.

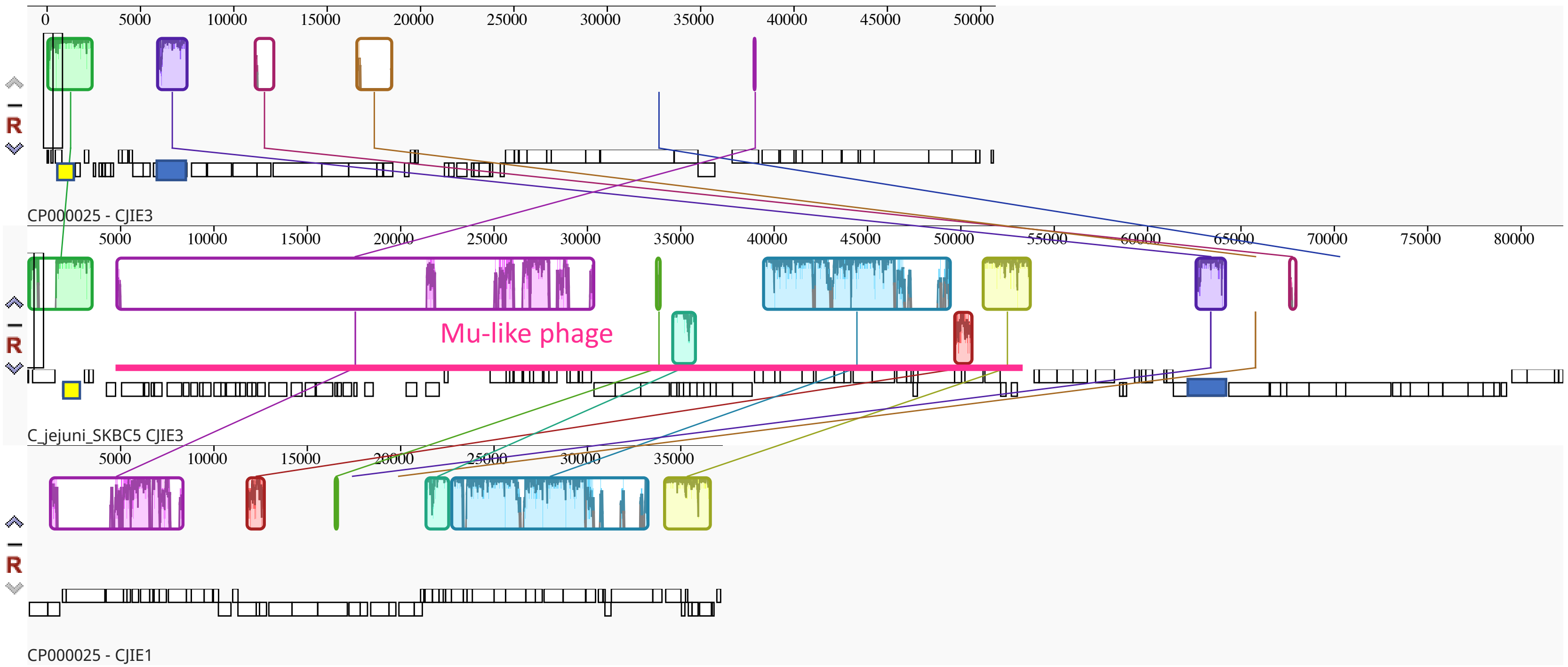


**Supplementary Figure 4. Comparison of element at CJIE3 insertion site.** Insertions and deletions were compared between CJIE3 in C. jejuni strain RM1221 and the element at the CJIE3 integration site at the tRNA-Arg gene near aroB and tgt in SKBC5. CJIE1 in C. jejuni strain RM1221 was included in the alignment with the order: RM1221 CJIE3, element in SKBC5 and RM1221 CJIE1. Alignment was created and visualized by Mauve software. Conserved blocks that were inverted compared to RM1221 in the figure are located beneath the SKBC5 integrated element. As mentioned, this element in SKBC5 possesses a Mu-like bacteriophage that is underlined with a pink bar. CJIE3 site specific-nuclease (integrase) and traG-like genes in both RM1221 and SKBC5 are colored yellow and blue, respectively.

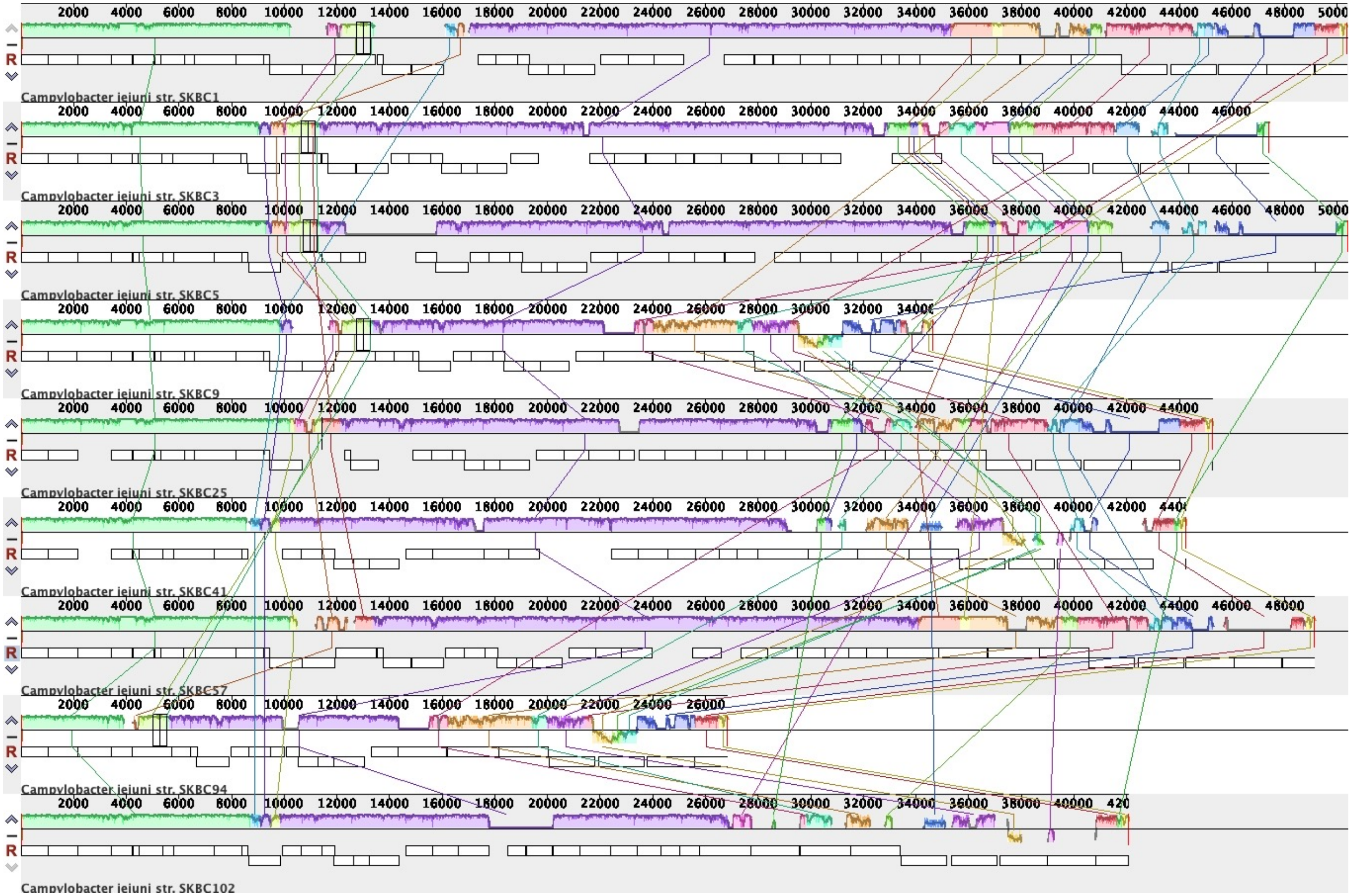


**Supplementary Figure 5. Comparisons of flagellar modification loci.** Flagellar modification (FM) biosynthesis loci alignment of the *C. jejuni* isolates from bears using Mauve revealed several local collinear blocks conserved among FM loci disrupted by insertions and deletions. The order of FM loci is SKBC1, SKBC3, SKBC5, SKBC9, SKBC25, SKBC41, SKBC57, SKBC94 and SKBC102. Conserved blocks that were inverted compared to SKBC1 or other previous strain in the figure are located beneath the FM loci.

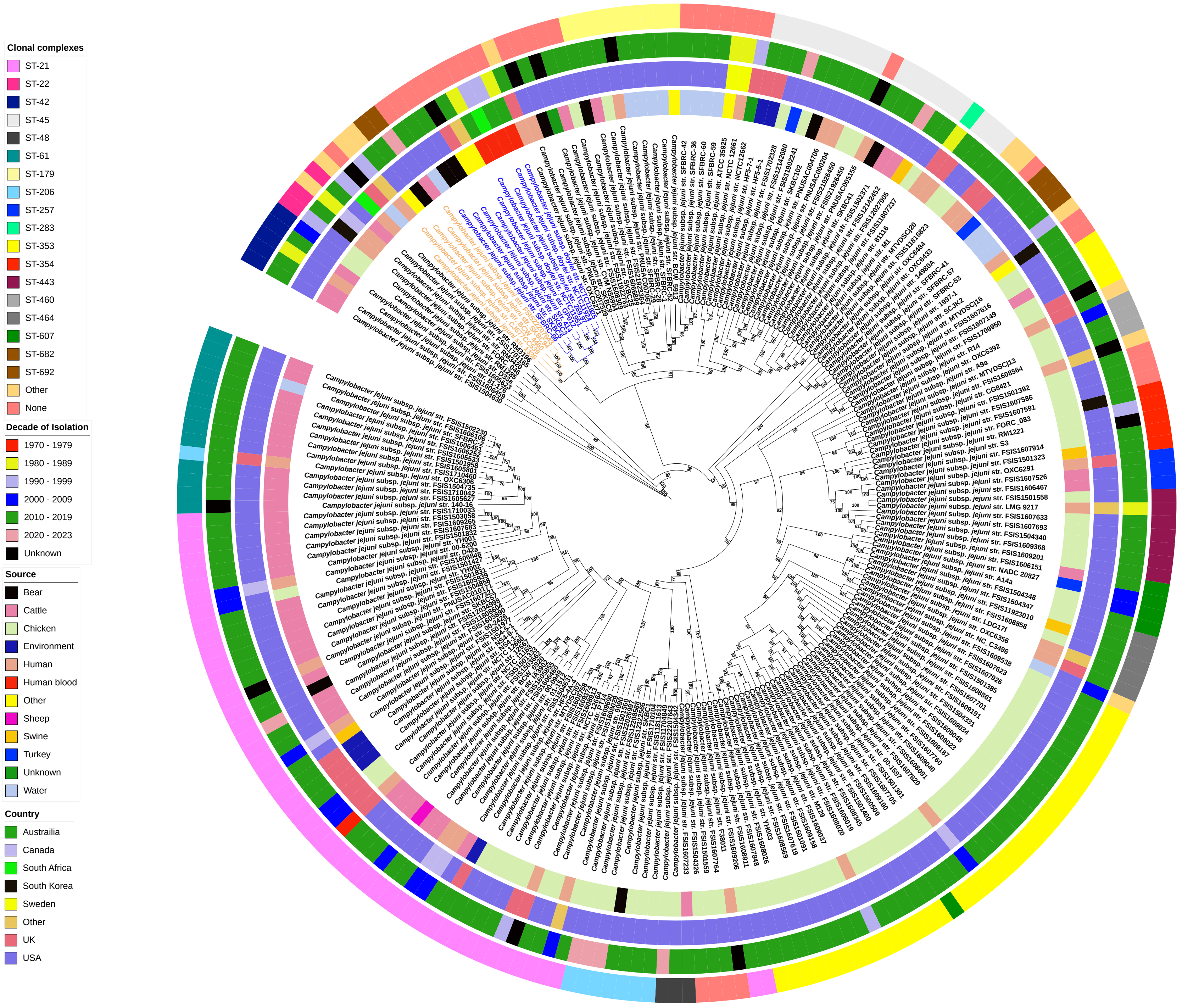


**Supplemental Figure 6.** *C. jejuni* isolates from bears (n=9) were compared with a global collection of publicly available genomes (n = 71) along with environmental and animal associated genomes isolated in the Southeastern U.S. (n=143). The 1,107 core genes from the genomes of all 223 genomic sequences were identified by Roary software and aligned using MAFFT. The dendrogram was constructed using RAxML with the GTRCAT model and 1,000 bootstraps. Clustered branches that are colored blue identify strains possessing deletions in both *cdt* and *mfr* loci, and those colored orange on adjacent branches possessing nonsense, point mutations in the *cdt* genes. Colored circles represent from inner to outer: (1) Source of isolation; (2) Country of isolation; (3) Decade of isolation; and (4) Clonal complex.
