## Supplementary Tables for "American black bear (*Ursus americanus*) as a potential host for *Campylobacter jejuni*"

**Supplementary Table 1: Strains used for comparison**

**GLOBAL**

| Strain | Taxon | source | location | Year | ST | CC | GenBank accession # |
|---|---|---|---|---|---|---|---|
| SKBC1 | *Cjj* | black bear feces | US: North Carolina | 2014 | 222 | 206 | CP125383 |
| SKBC102 | *Cjj* | black bear feces | US: North Carolina | 2016 | 45 | 45 | CP125387 |
| SKBC25 | *Cjj* | black bear feces | US: North Carolina | 2015 | 10624 | 179 | CP125388 |
| SKBC3 | *Cjj* | black bear feces | US: North Carolina | 2014 | 7630 | n/a | CP125394 |
| SKBC41 | *Cjj* | black bear feces | US: Virginia | 2015 | 45 | 45 | CP125386 |
| SKBC5 | *Cjj* | black bear feces | US: North Carolina | 2014 | 10620 | n/a | CP125391 |
| SKBC57 | *Cjj* | black bear feces | US: North Carolina | 2016 | 21 | 21 | CP125396 |
| SKBC9 | *Cjj* | black bear feces | US: Georgia | 2014 | 10501 | n/a | CP125390 |
| SKBC94 | *Cjj* | black bear feces | US: North Carolina | 2016 | 682 | 682 | CP125395 |
| LMG 9217 | *Cjj* | human stool | Belgium | 1986 | 443 | 443 | AIOO010000000 |
| 1997-1 | *Cjj* | human stool | USA | 1997 | 658 | 658 | AIOT010000000 |
| 140-16 | *Cjj* | cattle, stool | USA | Unk. | 5161 | 61 | AIPF01000000 |
| NCTC 11168 | *Cjj* | human stool | UK: England | Unk. | 43 | 21 | AL111168 |
| 1336 | *Cjj* | water | Unknown | Unk. | 841 | n/a | CM000854 |
| 414 | *Cjj* | bank vole | Unknown | Unk. | 3704 | n/a | CM000855 |
| RM1221 | *Cjj* | chicken meat | USA: California | 1997 | 354 | 354 | CP000025 |
| 81-176 | *Cjj* | human stool | US: Minnesota | 1982 | 604 | 42 | CP000538 |
| 81116 | *Cjj* | human stool | UK: England | 1981 | 267 | 283 | CP000814 |
| IA3902 | *Cjj* | sheep, aborted placenta | US: Iowa | 2009 | 8 | 21 | CP001876 |
| M1 | *Cjj* | human stool | UK: England | 2010 | 137 | 45 | CP001900 |
| S3 | *Cjj* | Unknown | Unknown | Unk. | 354 | 354 | CP001960 |
| PT14 | *Cjj* | human stool | UK: England | Unk. | 50 | 21 | CP003871 |
| R14 | *Cjj* | chicken | UK: England | Unk. | 356 | 353 | CP005081 |
| CG8421 | *Cjj* | human stool | Thailand: Bangkok | Unk. | 1919 | 52 | CP005388 |
| 00-2426 | *Cjj* | human stool | Canada | 2000 | 21 | 21 | CP006708 |
| F38011 | *Cjj* | human stool | Unknown | Unk. | 3644 | n/a | CP006851 |
| M129 | *Cjj* | human stool | Unknown | 1990 | 353 | 353 | CP007749 |
| D42a | *Cjj* | chicken cecum | USA | 2007 | 21 | 21 | CP007751 |
| MTVDSCj20 | *Cjj* | chicken cecum | US: Michigan | 2013 | 8785 | 45 | CP008787 |
| YH001 | *Cjj* | beef liver | US: Pennsylvania | 2014 | 806 | 21 | CP010058 |

| | | | | | | | |
|---|---|---|---|---|---|---|---|
| 01-1512 | *Cjj* | human stool | Canada | 2001 | 8 | 21 | CP010072 |
| 00-0949 | *Cjj* | human stool | Canada | 2000 | 8 | 21 | CP010301 |
| 00-1597 | *Cjj* | human stool | Canada | 2000 | 930 | n/a | CP010306 |
| 00-6200 | *Cjj* | human stool | Canada | 2000 | 806 | 21 | CP010307 |
| RM3196 | *Cjj* | human stool | South Africa: Capetown | 1996 | 362 | 362 | CP012690 |
| RM1285 | *Cjj* | chicken meat | USA: California | 1997 | 22 | 22 | CP015209 |
| 14980A | *Cjj* | turkey feces | US: North Carolina | 2014 | 1839 | n/a | CP017029 |
| MTVDSCj07 | *Cjj* | chicken cecum | US: Michigan | 2013 | 8789 | 21 | CP017031 |
| MTVDSCj13 | *Cjj* | chicken cecum | US: Michigan | 2013 | 460 | 460 | CP017032 |
| MTVDSCj16 | *Cjj* | chicken cecum | US: Michigan | 2013 | 1911 | n/a | CP017033 |
| RM3420 | *Cjj* | human stool | Canada | 1981 | 41 | 41 | CP017456 |
| ATCC 35925 | *Cjj* | pigeon | Sweden | 1984 | 5843 | n/a | CP020045 |
| LDG17f | *Cjj* | human stool | Czechia | 2016 | 464 | 464 | CP040015 |
| 9090 | *Cjj* | human stool | Slovenia | 2009 | 50 | 21 | CP040016 |
| NADC 20827 | *Cjj* | turkey feces | US: Iowa | 2005 | 1212 | 607 | CP045048 |
| NCTC 11951$^T$ | *Cjd* | human blood | Unknown | 1986 | 62 | n/a | LR134359 |
| 269.97 | *Cjd* | human blood | South Africa: Capetown | 1997 | 1845 | n/a | CP000768 |
| NCTC 11924 | *Cjd* | human blood | Unknown | 1999 | - | n/a | LR134530 |
| NCTC 11925 | *Cjd* | human blood | Australia | 1988 | 8767 | n/a | LS483295 |
| HF5-4A-4 | *Cj* | farm environment | UK | 2012 | 861 | 21 | CP007188 |
| HF5-5-1 | *Cj* | farm environment | UK | 2012 | 45 | 45 | CP007189 |
| HF5-7-1 | *Cj* | farm environment | UK | 2012 | 45 | 45 | CP007190 |
| NS4-1-1 | *Cj* | farm environment | UK | 2012 | 21 | 21 | CP007191 |
| NS4-5-1 | *Cj* | farm environment | UK | 2012 | 21 | 21 | CP007192 |
| NS4-9-1 | *Cj* | farm environment | UK | 2012 | 21 | 21 | CP007193 |
| CJ677CC539 | *Cj* | human stool | Finland | 1996 | 794 | 677 | CP010457 |
| FORC_046 | *Cj* | human stool | South Korea: Seoul | 2016 | 22 | 22 | CP017229 |
| BCW_6920 | *Cj* | cow, abortion | Unknown | 2011 | 8 | 21 | CP017673 |
| NCTC 12662 | *Cj* | Unknown | UK | 1992 | 5843 | n/a | CP019965 |
| YH002 | *Cj* | calf liver | US: Pennsylvania | 2014 | 982 | 21 | CP020776 |
| 12567 | *Cj* | chicken | UK: England | 2005 | 53 | 21 | CP028909 |
| NCTC 12660 | *Cj* | chicken | UK: England | 2005 | 21 | 21 | CP028910 |
| NCTC 12661 | *Cj* | human stool | Sweden | 1985 | 5843 | n/a | CP028911 |
| NCTC 12664 | *Cj* | chicken | UK: England | 1992 | 50 | 21 | CP028912 |
| FORC_083 | *Cj* | chicken meat | South Korea: Seoul | 2017 | 6849 | 354 | CP028933 |

| | | | | | | | |
|---|---|---|---|---|---|---|---|
| SCJK2 | *Cj* | mouse | South Korea: Gangwon | 2017 | 8388 | n/a | CP038862 |
| YH003 | *Cj* | chicken meat | US: Pennsylvania | 2014 | 353 | 353 | CP041584 |
| D33a | *Cj* | chicken cecum | US: Arizona | 2003 | 459 | 42 | CP058293 |
| A14a | *Cj* | chicken cecum | US: Kansas | 2003 | 1212 | 607 | CP058295 |
| A9a | *Cj* | chicken cecum | US: Kansas | 2003 | 2827 | 460 | CP058299 |
| OXC6271 | *Cj* | human stool | UK: England | 2011 | 508 | 508 | CUID01000000 |
| OXC6291 | *Cj* | human stool | UK: England | 2011 | 2030 | 257 | CUIW01000000 |
| OXC6306 | *Cj* | human stool | UK: England | 2011 | 273 | 206 | CUJO01000000 |
| OXC6356 | *Cj* | human stool | UK: England | 2011 | 464 | 464 | CULI01000001 |
| OXC6392 | *Cj* | human stool | UK: England | 2011 | 574 | 574 | CUMV01000001 |
| OXC6433 | *Cj* | human stool | UK: England | 2011 | 573 | 573 | CUOL01000001 |
| OXC6484 | *Cj* | human stool | UK: England | 2011 | 403 | 403 | CUQC01000001 |
| GP012 | *Cj* | guinea pig stool | Peru: Iquitos | 2019 | 10316 | n/a | JACRSF000000000 |
| BCW_4460 | *Cj* | rhesus macaque feces | USA: California | 2015 | 9259 | 177 | MKAJ01000001 |
| BCW_5913 | *Cj* | rhesus macaque feces | USA: California | 2016 | 6629 | 179 | MKES01000001 |

**GEOGRAPHICALLY LOCAL (Eastern USA; especially Georgia (GA), North Carolina (NC), Virginia(VA)):**

| **Isolate** | **Source** | **State** | **Year** | **ST** | **CC** | **PubMLST id** | **NCBI Biosample/SRA_accession #** |
|---|---|---|---|---|---|---|---|
| PNUSAC007067 | human stool | n/a | n/a | 2524 | 179 | 89537 | n/a |
| PNUSAC007529 | human stool | n/a | n/a | 10501 | n/a | 89602 | n/a |
| FSIS11811613 | chicken offal or meat | VA | 2018 | 222 | 206 | 91623 | n/a |
| FSIS1702328 | chicken offal or meat | GA | 2017 | 45 | 45 | 92249 | SAMN07410381; SRS2380429 |
| FSIS1609690 | chicken offal or meat | NC | 2016 | 50 | 21 | 92472 | SAMN06256283; SRS1938321 |
| FSIS1607586 | chicken offal or meat | GA | 2016 | 2083 | n/a | 92497 | SAMN05763277; SRS1689518 |
| FSIS1609037 | chicken offal or meat | GA | 2016 | 353 | 353 | 92504 | SAMN06140436; SRS1858539 |
| FSIS1607764 | chicken offal or meat | GA | 2016 | 6091 | n/a | 92518 | SAMN05833071; SRS1717570 |
| FSIS1607591 | chicken offal or meat | GA | 2016 | 2083 | n/a | 92530 | SAMN05799409; SRS1707549 |
| FSIS1607633 | chicken offal or meat | GA | 2016 | 51 | 443 | 92535 | SAMN05799436; SRS1707552 |
| FSIS11704849 | chicken offal or meat | GA | 2017 | 222 | 206 | 92624 | SAMN08224441; SRS2780950 |
| FSIS1710104 | chicken | GA | 2017 | 222 | 206 | 92662 | SAMN06328364; SRS1974250 |
| FSIS1609201 | chicken | GA | 2016 | 51 | 443 | 92695 | SAMN06179997; SRS1879691 |
| FSIS1608564 | chicken | GA | 2016 | 460 | 460 | 92734 | SAMN06046130; SRS1813091 |

| | | | | | | | |
|---|---|---|---|---|---|---|---|
| FSIS1608026 | chicken offal or meat | GA | 2016 | 353 | 353 | 92744 | SAMN05945123; SRS1761697 |
| FSIS1609374 | chicken offal or meat | GA | 2016 | 50 | 21 | 92750 | SAMN06210480; SRS1902216 |
| FSIS1609034 | chicken offal or meat | NC | 2016 | 3510 | 353 | 92764 | SAMN06140433; SRS1858531 |
| FSIS1608861 | chicken offal or meat | VA | 2016 | 353 | 353 | 92770 | SAMN06127095; SRS1847025 |
| FSIS1608913 | chicken offal or meat | GA | 2016 | 50 | 21 | 92795 | SAMN06127104; SRS1847027 |
| FSIS1608569 | chicken offal or meat | VA | 2016 | 353 | 353 | 92810 | SAMN06046135; SRS1813101 |
| FSIS1608023 | chicken | VA | 2016 | 3510 | 353 | 92820 | SAMN05945120; SRS1761716 |
| FSIS1501392 | chicken | NC | 2015 | 2083 | | 92873 | SAMN03785200; SRS969568 |
| FSIS1501558 | chicken offal or meat | GA | 2015 | 443 | 443 | 92876 | SAMN03850816; SRS985870 |
| FSIS1501391 | chicken offal or meat | GA | 2015 | 939 | 353 | 92878 | SAMN03785199; SRS969566 |
| FSIS1609045 | chicken offal or meat | GA | 2016 | 404 | 353 | 93114 | SAMN06140444; SRS1858547 |
| FSIS1608758 | chicken | GA | 2016 | 50 | 21 | 93123 | SAMN06127133; SRS1847102 |
| FSIS1608911 | chicken offal or meat | GA | 2016 | 454 | 21 | 93129 | SAMN06127102; SRS1847040 |
| FSIS1608858 | chicken offal or meat | GA | 2016 | 464 | 464 | 93132 | SAMN06127092; SRS1847029 |
| FSIS1608345 | chicken offal or meat | GA | 2016 | 939 | 353 | 93154 | SAMN06015823; SRS1797969 |
| FSIS1608019 | chicken | NC | 2016 | 353 | 353 | 93170 | SAMN05945116; SRS1761704 |
| FSIS1608020 | chicken offal or meat | VA | 2016 | 3515 | 353 | 93175 | SAMN05945117; SRS1761722 |
| FSIS1607914 | chicken offal or meat | GA | 2016 | 354 | 354 | 93176 | SAMN05928212; SRS1753175 |
| FSIS1607616 | chicken | NC | 2016 | 3736 | 353 | 93183 | SAMN05799424; SRS1707551 |
| FSIS1607623 | chicken offal or meat | GA | 2016 | 4370 | 353 | 93186 | SAMN05799430; SRS1707521 |
| FSIS1501400 | chicken offal or meat | GA | 2015 | 939 | 353 | 93200 | SAMN03785205; SRS969573 |
| FSIS1504340 | chicken | GA | 2015 | 51 | 443 | 93345 | SAMN04267339; SRS1162925 |
| FSIS1500509 | chicken | GA | 2013 | 939 | 353 | 93346 | SAMN03897488; SRS1010270 |
| FSIS1504347 | chicken offal or meat | VA | 2015 | 464 | 464 | 93348 | SAMN04267345; SRS1162923 |
| FSIS1501385 | chicken offal or meat | GA | 2015 | 353 | 353 | 93350 | SAMN03785194; SRS969558 |
| FSIS1609206 | chicken offal or meat | GA | 2016 | 454 | 21 | 93657 | SAMN06180002; SRS1879683 |
| FSIS1609368 | chicken offal or meat | GA | 2016 | 51 | 443 | 93659 | SAMN06256270; SRS1938334 |
| FSIS1709950 | chicken offal or meat | NC | 2016 | 3736 | 353 | 93686 | SAMN06276939; SRS1945030 |
| FSIS1609187 | chicken offal or meat | GA | 2016 | 10703 | 353 | 93690 | SAMN06179983; SRS1879695 |
| FSIS1609191 | chicken offal or meat | VA | 2016 | 353 | 353 | 93695 | SAMN06179987; SRS1879692 |
| FSIS1609538 | chicken offal or meat | NC | 2016 | 4370 | 353 | 93944 | SAMN06602732; SRS2049484 |
| FSIS1609040 | chicken offal or meat | VA | 2016 | 3510 | 353 | 93958 | SAMN06140439; SRS1858554 |
| FSIS1609190 | chicken offal or meat | GA | 2016 | 10751 | 607 | 93959 | SAMN06179986; SRS1879698 |
| FSIS1608029 | chicken offal or meat | GA | 2016 | 50 | 21 | 93961 | SAMN05945126; SRS1761711 |
| FSIS1608091 | chicken offal or meat | GA | 2016 | 3510 | 353 | 93962 | SAMN05945140; SRS1761712 |

| | | | | | | | |
|---|---|---|---|---|---|---|---|
| FSIS1607926 | chicken | VA | 2016 | 4370 | 353 | 93968 | SAMN05928223; SRS1753158 |
| FSIS1607848 | chicken offal or meat | GA | 2016 | 353 | 353 | 93969 | SAMN05900945; SRS1742329 |
| FSIS1607760 | chicken offal or meat | GA | 2016 | 4376 | 353 | 93977 | SAMN05833067; SRS1717579 |
| FSIS1607701 | chicken offal or meat | VA | 2016 | 353 | 353 | 93984 | SAMN05833099; SRS1717486 |
| FSIS1607705 | chicken offal or meat | GA | 2016 | 939 | 353 | 93985 | SAMN05833103; SRS1717495 |
| FSIS1607620 | chicken offal or meat | GA | 2016 | 2132 | 353 | 93988 | SAMN05799427; SRS1707438 |
| FSIS1607619 | chicken offal or meat | NC | 2016 | 10520 | 353 | 93990 | SAMN05799426; SRS1707447 |
| FSIS1504326 | chicken offal or meat | GA | 2015 | 6091 | n/a | 94009 | SAMN04267325; SRS1162916 |
| FSIS1504342 | chicken offal or meat | NC | 2015 | 3694 | 48 | 94012 | SAMN04267341; SRS1162920 |
| FSIS1504348 | chicken offal or meat | NC | 2015 | 1212 | 607 | 94017 | SAMN04267346; SRS1163359 |
| FSIS1504331 | chicken offal or meat | GA | 2015 | 10221 | 353 | 94021 | SAMN04267330; SRS1163509 |
| FSIS1501091 | chicken offal or meat | NC | 2015 | 2829 | 353 | 94028 | SAMN03795224; SRS973092 |
| FSIS1501559 | chicken offal or meat | GA | 2015 | 6091 | n/a | 94033 | SAMN03850817; SRS985871 |
| FSIS1501560 | chicken offal or meat | GA | 2015 | 50 | 21 | 94037 | SAMN03850818; SRS985873 |
| FSIS11816823 | chicken | GA | 2018 | 137 | 45 | 94752 | SAMN10881670; SRS4337209 |
| FSIS21923364 | chicken offal or meat | NY | 2019 | 2524 | 179 | 94767 | SAMN10880123; SRS4333512 |
| CVM N55904 | chicken offal or meat | MO | 2017 | 10501 | n/a | 97941 | SAMN06660919; SRS2099146 |
| FSIS1606899 | cattle | TN | 2016 | 2524 | 179 | 98512 | SAMN05366799, SRS1548480 |
| FSIS1607233 | cattle | NC | 2016 | 918 | 48 | 99391 | SAMN05558641; SRS1609336 |
| FSIS1607149 | cattle | NC | 2016 | 3736 | 353 | 99419 | SAMN05505603; SRS1598342 |
| FSIS1606848 | cattle | NC | 2016 | 982 | 21 | 99559 | SAMN05301270; SRS1529695 |
| FSIS1606461 | cattle | GA | 2016 | 61 | 61 | 99598 | SAMN05190020; SRS1475011 |
| FSIS1606460 | cattle | GA | 2016 | 8 | 21 | 99604 | SAMN05190019; SRS1475009 |
| FSIS1606459 | cattle | VA | 2016 | 42 | 42 | 99607 | SAMN05190018; SRS1475008 |
| FSIS1606467 | cattle | VA | 2016 | 6673 | 257 | 99673 | SAMN05190026; SRS1475017 |
| FSIS1606252 | cattle | GA | 2016 | 61 | 61 | 99810 | SAMN04625697; SRS1382924 |
| FSIS1605855 | cattle | GA | 2016 | 8 | 21 | 99833 | SAMN04524182; SRS1316544 |
| FSIS1606151 | cattle | NC | 2016 | 607 | 607 | 99840 | SAMN04600084; SRS1371155 |
| FSIS1606106 | cattle | NC | 2016 | 61 | 61 | 99842 | SAMN04600066; SRS1371151 |
| FSIS1605890 | cattle | NC | 2016 | 21 | 21 | 99904 | SAMN04526075; SRS1318878 |
| FSIS1605939 | cattle | NC | 2016 | 982 | 21 | 99928 | SAMN04537006; SRS1327433 |
| FSIS1605623 | cattle | NC | 2016 | 459 | 42 | 100060 | SAMN04461472; SRS1278043 |
| FSIS1605801 | cattle | GA | 2016 | 61 | 61 | 100061 | SAMN04508250; SRS1307320 |
| FSIS1605627 | cattle | NC | 2016 | 1244 | 61 | 100074 | SAMN04461490; SRS1277875 |
| FSIS1607321 | cattle | GA | 2016 | 21 | 21 | 100701 | SAMN05578358; SRS1620915 |

| | | | | | | | |
|---|---|---|---|---|---|---|---|
| FSIS11807237 | chicken | VA | 2017 | 45 | 45 | 101041 | SAMN08437409; SRS2898277 |
| FSIS1607526 | cattle | NC | 2016 | 929 | 257 | 101272 | SAMN05763253; SRS1689521 |
| FSIS1609265 | cattle | VA | 2016 | 806 | 21 | 101332 | SAMN06179687; SRS1878018 |
| FSIS1504735 | cattle | GA | 2015 | 1244 | 61 | 101494 | SAMN04452114; SRS1272300 |
| FSIS1502230 | cattle | NC | 2015 | 61 | 61 | 101498 | SAMN04388419; SRS1240223 |
| FSIS1503662 | cattle | GA | 2015 | 8 | 21 | 101507 | SAMN04388448; SRS1240204 |
| FSIS1501831 | cattle | NC | 2015 | 982 | 21 | 101538 | SAMN04320879; SRS1192310 |
| FSIS1501958 | cattle | NC | 2015 | 61 | 61 | 101539 | SAMN04320886; SRS1192366 |
| FSIS1504630 | cattle | NC | 2015 | 42 | 42 | 101554 | SAMN04195060; SRS1121793 |
| FSIS1501333 | cattle | NC | 2015 | 8 | 21 | 101567 | SAMN04091194; SRS1071596 |
| FSIS1609158 | chicken | GA | 2016 | 353 | 353 | 101870 | SAMN06165733; SRS1871244 |
| FSIS1605533 | cattle | NC | 2015 | 61 | 61 | 102020 | SAMN04444345; SRS1266750 |
| FSIS1503058 | cattle | GA | 2015 | 806 | 21 | 102021 | SAMN04388427; SRS1240219 |
| FSIS1504631 | cattle | NC | 2015 | 21 | 21 | 102067 | SAMN04195061; SRS1121792 |
| FSIS1501427 | cattle | GA | 2015 | 982 | 21 | 102223 | SAMN04027116; SRS1054389 |
| FSIS1710033 | cattle | GA | 2016 | 806 | 21 | 103340 | SAMN06276449; SRS1945037 |
| FSIS1710042 | cattle | NC | 2016 | 1244 | 61 | 103348 | SAMN06276453; SRS1945001 |
| FSIS1701165 | cattle | GA | 2017 | 22 | 22 | 103991 | SAMN06915353; SRS2172035 |
| FSIS1710460 | cattle | GA | 2016 | 61 | 61 | 104131 | SAMN06346617; SRS1991268 |
| FSIS1608309 | cattle | TN | 2016 | 10501 | | 104166 | SAMN06046063; SRS1812533 |
| FSIS1607683 | cattle | VA | 2016 | 806 | 21 | 104208 | SAMN05833081; SRS1717485 |
| FSIS1607693 | chicken | GA | 2016 | 51 | 443 | 104218 | SAMN05833091; SRS1717476 |
| FSIS1501977 | swine | VA | 2015 | 21 | 21 | 104332 | SAMN04320901; SRS1192377 |
| FSIS1501832 | cattle | NC | 2015 | 806 | 21 | 104333 | SAMN04320880; SRS1192304 |
| FSIS1504858 | cattle | NC | 2015 | 982 | 21 | 104343 | SAMN04229854; SRS1144016 |
| FSIS1501323 | swine | NC | 2015 | 13296 | 354 | 104370 | SAMN04091185; SRS1071587 |
| SFBRC-1 | environmental water | GA | 2012 | 3889 | 179 | 106318 | n/a |
| SFBRC-2 | environmental water | GA | 2012 | 61 | 61 | 106319 | n/a |
| SFBRC-28 | environmental water | GA | 2013 | 2524 | 179 | 106336 | n/a |
| SFBRC-29 | environmental water | GA | 2013 | 2524 | 179 | 106337 | n/a |
| SFBRC-36 | environmental water | GA | 2013 | 7943 | | 106344 | n/a |
| SFBRC-41 | environmental water | GA | 2013 | 699 | 692 | 106347 | n/a |
| SFBRC-42 | environmental water | GA | 2013 | 2866 | | 106348 | n/a |
| SFBRC-52 | environmental water | GA | 2013 | 3889 | 179 | 106350 | n/a |

| | | | | | | | |
|---|---|---|---|---|---|---|---|
| SFBRC-53 | environmental water | GA | 2013 | 692 | 692 | 106351 | n/a |
| SFBRC-57 | environmental water | GA | 2013 | 692 | 692 | 106352 | n/a |
| SFBRC-59 | environmental water | GA | 2013 | 7943 | n/a | 106353 | n/a |
| SFBRC-60 | environmental water | GA | 2013 | 7949 | n/a | 106354 | n/a |
| SFBRC-66 | environmental water | GA | 2013 | 7949 | n/a | 106355 | n/a |
| FSIS11922763 | chicken | KY | 2019 | 10501 | n/a | 120341 | SAMN12349472; SRS5145307 |
| FSIS11923010 | swine | VA | 2019 | 464 | 464 | 120435 | SAMN12345917; SRS5144437 |
| FSIS12027905 | swine | NC | 2019 | 45 | 45 | 122671 | SAMN14737992; SRS6540860 |
| FSIS12106873 | cattle | PA | 2021 | 682 | 682 | 126827 | SAMN23493042; SRS11178343 |
| FSIS12142080 | turkey | VA | 2021 | 45 | 45 | 128966 | SAMN21463028; SRS10189879 |
| FSIS12142452 | cattle | NC | 2021 | 45 | 45 | 129106 | SAMN21337215; SRS10053674 |
| FSIS21926450 | chicken offal or meat | GA | 2019 | 45 | 45 | 134474 | SAMN13708426; SRS5931241 |
| FSIS22207643 | chicken offal or meat | VA | 2022 | 48 | 48 | 138360 | SAMN27522664; SRS12574200 |
| FSIS11808604 | cattle | NC | 2018 | 21 | 21 | 102857 | SAMN08816081; SRS3106598 |
| FSIS12034998 | cattle | NC | 2020 | 21 | 21 | 125437 | SAMN16811271; SRS7723740 |
| FSIS12208588 | chicken carcass | NC | 2021 | 222 | 206 | 129596 | SAMN25159901; SRS11733377 |
| FSIS12322565 | raw intact chicken | AL | 2023 | n/a | n/a | n/a | SAMN35685631; SRS17944688 |
| FSIS1502371 | cattle | NJ | 2015 | 45 | 45 | 101541 | SAMN04120349; SRS1181114 |
| PNUSAC000204 | human stool | n/a | n/a | 6647 | 49 | 85526 | SAMN04339734; SRS1211470 |
| PNUSAC004706 | human stool | n/a | n/a | 10446 | 45 | 82515 | SAMN09292332; SRS3363973 |
| PNUSAC005155 | human stool | n/a | n/a | 45 | 45 | 81938 | SAMN09665077; SRS3549084 |
| PNUSAC010171 | human stool | n/a | n/a | n/a | n/a | n/a | SAMN12585128; SRS5273649 |
| FSIS22026997 | raw intact chicken | GA | 2020 | 222 | 206 | 134728 | SAMN14085345; SRS6133342 |
| FSIS31902241 | chicken carcass | GA | 2019 | 45 | 45 | 140604 | SAMN12163344; SRS5043817 |
| NC_C3496 | water | NC | 2009 | n/a | n/a | n/a | SAMN08168977; SRS3211689 |

### Supplementary Table 2: Strains in PubMLST that share four alleles with either SKBC9 or SKBC25

| Strain | Location[a] | Year | source | *aspA* | *glnA* | *gltA* | *glyA* | *pgm* | *tkt* | *uncA* | ST | CC[b] |
|---|---|---|---|---|---|---|---|---|---|---|---|---|
| 06-1507 | Canada:Quebec | 2006 | env. water | 1 | 6 | 61 | 244 | 40 | 32 | 3 | 2524 | 179 |
| 007A-0395 | Canada:Ontario | 2005 | env. water | 1 | 6 | 61 | 176 | 40 | 32 | 3 | 4380 | 179 |
| BICO 301 (14) | USA:Georgia | 2005 | env. water | 1 | 6 | 61 | 244 | 40 | 32 | 3 | 2524 | 179 |
| NORO 513 (83a) | USA:Georgia | 2007 | env. water | 1 | 6 | 61 | 244 | 40 | 32 | 3 | 2524 | 179 |
| NORO 115 (38a) | USA:Georgia | 2008 | env. water | 1 | 6 | 61 | 244 | 40 | 32 | 3 | 2524 | 179 |
| SFBRC-12 | USA:Georgia | 2012 | env. water | 1 | 6 | 61 | 244 | 40 | 32 | 3 | 2524 | 179 |
| SFBRC-48 | USA:Georgia | 2013 | env. water | 1 | 6 | 61 | 244 | 40 | 32 | 3 | 2524 | 179 |
| PNUSAC009587 | USA | Unk. | human | 1 | 6 | 61 | 244 | 40 | 32 | 3 | 2524 | 179 |
| PNUSAC009549 | USA | Unk. | human | 1 | 6 | 61 | 244 | 40 | 32 | 3 | 2524 | 179 |
| PNUSAC001967 | USA:HHS_region 2 | 2016 | human | 1 | 6 | 61 | 244 | 40 | 32 | 3 | 2524 | 179 |
| PNUSAC005720 | USA:HHS_region 5 | 2018 | human | 1 | 6 | 61 | 176 | 40 | 32 | 3 | 4380 | 179 |
| PNUSAC005568 | USA:HHS_region 4 | 2018 | human | 1 | 6 | 61 | 244 | 40 | 32 | 3 | 2524 | 179 |
| PNUSAC005579 | USA:HHS_region 3 | 2018 | human | 1 | 6 | 61 | 244 | 40 | 32 | 3 | 2524 | 179 |
| PNUSAC005116 | USA:HHS_region 3 | 2018 | human | 1 | 6 | 61 | 244 | 40 | 32 | 3 | 2524 | 179 |
| PNUSAC009231 | USA | Unk. | human | 1 | 6 | 61 | 244 | 40 | 32 | 3 | 2524 | 179 |
| PNUSAC002170 | USA:HHS_region 2 | 2016 | human | 1 | 6 | 61 | 244 | 40 | 32 | 3 | 2524 | 179 |
| PNUSAC003437 | USA:HHS_region 5 | 2017 | human | 12 | 6 | 61 | 147 | 261 | 32 | 3 | 10501 | NA |
| PNUSAC004034 | USA:Wyoming | 2011 | cattle | 1 | 6 | 61 | 244 | 40 | 32 | 3 | 2524 | 179 |
| PNUSAC001646 | USA:HHS_region 1 | 2017 | human | 1 | 6 | 61 | 176 | 40 | 32 | 3 | 4380 | 179 |
| PNUSAC001432 | USA:HHS_region 3 | 2016 | human | 12 | 6 | 61 | 244 | 40 | 32 | 3 | 10537 | 179 |
| PNUSAC004020 | USA:HHS_region 5 | 2017 | human | 1 | 6 | 61 | 176 | 40 | 32 | 3 | 4380 | 179 |
| PNUSAC003948 | USA:HHS_region 4 | 2017 | human | 1 | 6 | 61 | 244 | 40 | 32 | 3 | 2524 | 179 |
| PNUSAC003780 | USA:HHS_region 3 | 2017 | human | 1 | 6 | 61 | 244 | 40 | 32 | 3 | 2524 | 179 |
| PNUSAC003495 | USA:HHS_region 3 | 2017 | human | 1 | 6 | 61 | 244 | 40 | 32 | 3 | 2524 | 179 |
| PNUSAC002060 | USA:HHS_region 8 | 2017 | human | 1 | 6 | 61 | 244 | 40 | 32 | 3 | 2524 | 179 |
| PNUSAC003351 | USA:HHS_region 3 | 2017 | human | 1 | 6 | 61 | 244 | 40 | 32 | 3 | 2524 | 179 |
| PNUSAC002955 | USA:HHS_region 5 | 2017 | human | 1 | 6 | 61 | 244 | 40 | 32 | 3 | 2524 | 179 |
| PNUSAC007552 | USA | Unk. | human | 1 | 6 | 61 | 244 | 40 | 32 | 3 | 2524 | 179 |
| PNUSAC007662 | USA | Unk. | human | 1 | 6 | 61 | 244 | 40 | 32 | 3 | 2524 | 179 |
| PNUSAC007820 | USA | Unk. | human | 1 | 6 | 61 | 244 | 40 | 32 | 3 | 2524 | 179 |
| PNUSAC008247 | USA | Unk. | human | 1 | 6 | 61 | 244 | 40 | 32 | 3 | 2524 | 179 |
| PNUSAC007856 | USA | Unk. | human | 1 | 6 | 61 | 244 | 40 | 32 | 3 | 2524 | 179 |
| PNUSAC007714 | USA | Unk. | human | 1 | 6 | 61 | 244 | 40 | 32 | 3 | 2524 | 179 |
| PNUSAC007690 | USA | Unk. | human | 1 | 6 | 61 | 244 | 40 | 32 | 3 | 2524 | 179 |

| | | | | | | | | | | | | |
|---|---|---|---|---|---|---|---|---|---|---|---|---|
| PNUSAC007067 | USA | Unk. | human | 1 | 6 | 61 | 244 | 40 | 32 | 3 | 2524 | 179 |
| PNUSAC007535 | USA | Unk. | human | 1 | 6 | 61 | 176 | 40 | 32 | 3 | 4380 | 179 |
| PNUSAC007529 | USA | Unk. | human | 12 | 6 | 61 | 147 | 261 | 32 | 3 | 10501 | NA |
| PNUSAC007290 | USA | Unk. | human | 12 | 6 | 61 | 147 | 261 | 32 | 3 | 10501 | NA |
| PNUSAC007217 | USA | Unk. | human | 1 | 6 | 61 | 244 | 40 | 32 | 3 | 2524 | 179 |
| PNUSAC007216 | USA | Unk. | human | 1 | 6 | 61 | 244 | 40 | 32 | 3 | 2524 | 179 |
| PNUSAC007165 | USA | Unk. | human | 1 | 6 | 61 | 244 | 40 | 32 | 3 | 2524 | 179 |
| PNUSAC007116 | USA | Unk. | human | 1 | 6 | 61 | 244 | 40 | 32 | 3 | 2524 | 179 |
| PNUSAC006882 | USA:HHS_region 5 | 2018 | human | 1 | 6 | 61 | 176 | 40 | 32 | 3 | 4380 | 179 |
| PNUSAC006826 | USA:HHS_region 1 | 2018 | human | 1 | 6 | 61 | 244 | 40 | 32 | 3 | 2524 | 179 |
| PNUSAC006211 | USA:HHS_region 3 | 2018 | human | 1 | 6 | 61 | 176 | 40 | 32 | 3 | 4380 | 179 |
| PNUSAC006032 | USA:HHS_region 5 | 2018 | human | 1 | 6 | 61 | 244 | 40 | 32 | 3 | 2524 | 179 |
| PNUSAC006234 | USA:HHS_region 4 | 2018 | human | 1 | 6 | 61 | 244 | 40 | 32 | 3 | 2524 | 179 |
| PNUSAC006588 | USA:HHS_region 4 | 2018 | human | 1 | 6 | 61 | 244 | 40 | 32 | 3 | 2524 | 179 |
| PNUSAC006687 | USA:HHS_region 5 | 2018 | human | 1 | 6 | 61 | 244 | 40 | 32 | 3 | 2524 | 179 |
| PNUSAC006589 | USA:HHS_region 5 | 2018 | human | 1 | 6 | 61 | 176 | 40 | 32 | 3 | 4380 | 179 |
| FSIS21720490 | USA:Michigan | 2017 | chicken | 1 | 6 | 61 | 244 | 40 | 32 | 3 | 2524 | 179 |
| FSIS21923364 | USA:New York | 2019 | chicken | 1 | 6 | 61 | 244 | 40 | 32 | 3 | 2524 | 179 |
| CVM N17C624 | USA:Texas | 2017 | chicken | 1 | 6 | 61 | 244 | 40 | 32 | 3 | 2524 | 179 |
| CVM N55904 | USA:Missouri | 2015 | chicken | 12 | 6 | 61 | 147 | 261 | 32 | 3 | 10501 | NA |
| CVM N18C002 | USA:California | 2018 | turkey | 1 | 6 | 61 | 176 | 40 | 32 | 3 | 4380 | 179 |
| FSIS1606899 | USA:Tennessee | 2016 | cattle | 1 | 6 | 61 | 244 | 40 | 32 | 3 | 2524 | 179 |
| FSIS11812606 | USA:Iowa | 2018 | chicken | 1 | 6 | 61 | 176 | 40 | 32 | 3 | 4380 | 179 |
| FSIS1608309 | USA:Tennessee | 2016 | cattle | 12 | 6 | 61 | 147 | 261 | 32 | 3 | 10501 | NA |
| **SKBC9** | **USA:Georgia** | **2014** | **black bear** | 12 | 6 | 61 | 147 | 261 | 32 | 3 | 10501 | NA |
| **SKBC25** | **USA:North Carolina** | **2015** | **black bear** | 1 | 6 | 61 | 244 | 797 | 32 | 3 | 10624 | 179 |
| SFBRC-28 | USA:Georgia | 2013 | env. water | 1 | 6 | 61 | 244 | 40 | 32 | 3 | 2524 | 179 |
| SFBRC-29 | USA:Georgia | 2013 | env. water | 1 | 6 | 61 | 244 | 40 | 32 | 3 | 2524 | 179 |
| SFBRC-32 | USA:Georgia | 2013 | env. water | 1 | 6 | 61 | 244 | 40 | 32 | 3 | 2524 | 179 |
| FSIS11922763 | USA:Kentucky | 2019 | chicken | 12 | 6 | 61 | 147 | 261 | 32 | 3 | 10501 | NA |
| FSIS11926407 | USA:Ohio | 2019 | turkey | 1 | 6 | 61 | 244 | 40 | 32 | 3 | 2524 | 179 |
| FSIS12028294 | USA:Arkansas | 2020 | chicken | 1 | 6 | 61 | 244 | 40 | 32 | 3 | 2524 | 179 |
| FSIS12036743 | USA:Pennsylvania | 2020 | turkey | 1 | 6 | 61 | 244 | 40 | 32 | 3 | 2524 | 179 |
| FSIS12107699 | USA:Arkansas | 2021 | chicken | 1 | 6 | 61 | 244 | 40 | 32 | 3 | 2524 | 179 |
| FSIS12214444 | USA:Iowa | 2022 | cattle | 1 | 6 | 61 | 176 | 40 | 32 | 3 | 4380 | 179 |
| FSIS12320378 | USA:Iowa | 2023 | chicken | 1 | 6 | 61 | 176 | 40 | 32 | 3 | 4380 | 179 |
| FSIS32206488 | USA:Arkansas | 2022 | chicken | 1 | 6 | 61 | 176 | 40 | 32 | 3 | 4380 | 179 |
| FSIS32206495 | USA:Arkansas | 2022 | chicken | 1 | 6 | 61 | 176 | 40 | 32 | 3 | 4380 | 179 |

[a]: HHS_region 1 = (Connecticut, Maine, Massachusetts, New Hampshire, Rhode Island, Vermont)
HHS_region 2 = (New Jersey, New York, Puerto Rico, U.S. Virgin Islands)
HHS_region 3 = (Delaware, Maryland, Pennsylvania, Virginia, West Virginia, Washington D.C.)
HHS_region 4 = (Alabama, Florida, Georgia, Kentucky, Mississippi, North Carolina, South Carolina, Tennessee)
HHS_region 5 = (Illinois, Indiana, Michigan, Minnesota, Ohio, Wisconsin)
HHS_region 8 = (Colorado, Montana, North Dakota, South Dakota, Utah, Wyoming)

[b]: NA, not assigned

### Supplementary Table 3: Strains in PubMLST with CC ST-682

| Strain | Location | year | source | *aspA* | *glnA* | *gltA* | *glyA* | *pgm* | *tkt* | *uncA* | ST |
|---|---|---|---|---|---|---|---|---|---|---|---|
| 78965 | UK | 1994 | sand (bathing beach) | 39 | 5 | 9 | 4 | 8 | 46 | 21 | 175 |
| 78970 | UK | 1994 | sand (bathing beach) | 37 | 2 | 9 | 2 | 8 | 46 | 23 | 176 |
| 87035 | UK | 1994 | sand (bathing beach) | 26 | 2 | 9 | 51 | 8 | 46 | 5 | 208 |
| 78966 | UK | 1994 | sand (bathing beach) | 2 | 2 | 9 | 51 | 8 | 46 | 5 | 215 |
| WB CL95004 c16.7.02 | UK | 2002 | starling | 26 | 2 | 9 | 51 | 8 | 46 | 21 | 682 |
| WB CL95011 c16.7.02 | UK | 2002 | starling | 17 | 2 | 9 | 51 | 121 | 46 | 21 | 686 |
| WB CL95012 c16.7.02 | UK | 2002 | starling | 26 | 43 | 9 | 101 | 121 | 46 | 21 | 687 |
| starling 4 | UK: Oxfordshire | 2002 | starling feces | 26 | 2 | 9 | 51 | 8 | 46 | 21 | 682 |
| starling 11 | UK: Oxfordshire | 2002 | starling feces | 17 | 2 | 9 | 51 | 121 | 46 | 21 | 686 |
| starling 12 | UK: Oxfordshire | 2002 | starling feces | 26 | 43 | 9 | 101 | 121 | 46 | 21 | 687 |
| WB CL95003 c16.7.02 | UK: Oxfordshire | 2002 | starling | 35 | 43 | 9 | 5 | 8 | 46 | 21 | 681 |
| WB CL95037e c6.2.03 | UK: Wytham | 2003 | starling | 35 | 2 | 8 | 51 | 121 | 46 | 21 | 818 |
| EX114 | UK | 1997 | broiler environment | 4 | 2 | 9 | 5 | 8 | 46 | 21 | 914 |
| CW04619 c15.6.04 | UK: Oxfordshire | 2004 | starling | 26 | 2 | 8 | 51 | 8 | 46 | 1 | 1019 |
| CW04620 c15.6.04 | UK: Oxfordshire | 2004 | starling | 26 | 43 | 9 | 101 | 8 | 46 | 21 | 1020 |
| WB CW04656 c15.6.04 | UK | 2004 | starling | 17 | 2 | 9 | 5 | 8 | 46 | 21 | 1021 |
| WB CW04658 c16.6.04 | UK: Oxfordshire | 2004 | starling | 26 | 43 | 9 | 51 | 121 | 46 | 21 | 1022 |
| ct86869 c4.6.04 | UK: Wytham | 2004 | starling | 26 | 2 | 9 | 51 | 8 | 2 | 21 | 1385 |
| ct86858 c4.6.04 | UK: Wytham | 2004 | starling | 26 | 2 | 9 | 101 | 8 | 46 | 21 | 1386 |
| ct86880 c8.6.04 | UK: Wytham | 2004 | starling | 35 | 43 | 9 | 51 | 8 | 2 | 21 | 1387 |
| cw04658 c27.6.04 | UK: Wytham | 2004 | starling | 26 | 43 | 9 | 51 | 8 | 46 | 21 | 1390 |
| ct43856 c26.1.04 | UK: Wytham | 2004 | starling | 26 | 43 | 9 | 2 | 8 | 46 | 21 | 1391 |
| cw04605 c25.6.04 | UK: Wytham | 2004 | starling | 26 | 2 | 9 | 5 | 8 | 46 | 21 | 1392 |
| cw04692 c22.6.04 | UK: Wytham | 2004 | starling | 26 | 2 | 9 | 51 | 121 | 46 | 21 | 1542 |
| ct86898 | UK | 2004 | starling | 26 | 43 | 9 | 5 | 121 | 46 | 21 | 1503 |
| CW04629 | UK | 2004 | starling | 17 | 2 | 8 | 51 | 121 | 46 | 21 | 1505 |
| CW04707 | UK | 2004 | starling | 17 | 5 | 9 | 101 | 8 | 46 | 21 | 1507 |
| cw04717 | UK | 2004 | starling | 26 | 2 | 9 | 51 | 121 | 46 | 21 | 1542 |
| 7423 | UK: Grampian | 2006 | wild bird | 35 | 2 | 9 | 51 | 8 | 2 | 21 | 1027 |
| WB CT43886 c26.1.4 | UK | 2004 | starling | 26 | 43 | 9 | 2 | 8 | 46 | 21 | 1391 |
| cl95027c c2.6.2003 | UK | 2003 | starling | 26 | 2 | 9 | 51 | 8 | 46 | 21 | 682 |
| WB CT43816 r11.12.03 | UK | 2003 | starling | 35 | 2 | 9 | 51 | 8 | 2 | 21 | 1027 |
| WB CL95027p c6.2.03 | UK | 2003 | starling | 26 | 2 | 9 | 51 | 8 | 46 | 21 | 682 |
| WB CT86845 c25.5.04 | UK | 2004 | starling | 26 | 43 | 9 | 101 | 8 | 46 | 21 | 1020 |

| | | | | | | | | | | | |
|---|---|---|---|---|---|---|---|---|---|---|---|
| WB CT86846 c25.5.04 | UK | 2004 | starling | 26 | 43 | 9 | 101 | 8 | 46 | 21 | 1020 |
| WB CT86848 c25.5.04 | UK | 2004 | starling | 26 | 2 | 9 | 51 | 8 | 46 | 21 | 682 |
| WB CT86849 c25.5.04 | UK | 2004 | starling | 26 | 2 | 9 | 51 | 8 | 46 | 21 | 682 |
| WB CT86851 c2.6.04 | UK | 2004 | starling | 26 | 43 | 9 | 101 | 8 | 46 | 21 | 1020 |
| WB CT86854 c2.6.04 | UK | 2004 | starling | 26 | 43 | 9 | 101 | 8 | 46 | 21 | 1020 |
| WB CL95083 c2.6.04 | UK | 2004 | starling | 26 | 43 | 9 | 101 | 8 | 46 | 21 | 1020 |
| WB CT86870 c4.6.04 | UK | 2004 | starling | 26 | 2 | 9 | 51 | 8 | 46 | 21 | 682 |
| WB CT86874 c4.6.04 | UK | 2004 | starling | 26 | 43 | 9 | 101 | 8 | 46 | 21 | 1020 |
| WB CL95044 c4.6.04 | UK | 2004 | starling | 26 | 43 | 9 | 101 | 8 | 46 | 21 | 1020 |
| WB CL95059 c4.6.04 | UK | 2004 | starling | 26 | 43 | 9 | 101 | 8 | 46 | 21 | 1020 |
| WB CL95083 c4.6.04 | UK | 2004 | starling | 26 | 43 | 9 | 101 | 8 | 46 | 21 | 1020 |
| WB CT86791 c4.6.04 | UK | 2004 | starling | 26 | 43 | 9 | 101 | 8 | 46 | 21 | 1020 |
| WB CT86854 c4.6.04 | UK | 2004 | starling | 26 | 43 | 9 | 101 | 8 | 46 | 21 | 1020 |
| WB CT86858 c4.6.04 | UK | 2004 | starling | 26 | 2 | 9 | 101 | 8 | 46 | 21 | 1386 |
| WB CT86878 c8.6.04 | UK | 2004 | starling | 35 | 2 | 9 | 51 | 8 | 2 | 21 | 1027 |
| WB CT86879 c8.6.04 | UK | 2004 | starling | 35 | 2 | 9 | 51 | 8 | 2 | 21 | 1027 |
| WB CT86882 c8.6.04 | UK | 2004 | starling | 26 | 43 | 9 | 101 | 8 | 46 | 21 | 1020 |
| WB CT86883 c8.6.04 | UK | 2004 | starling | 26 | 43 | 9 | 101 | 8 | 46 | 21 | 1020 |
| WB CT86888 c10.6.04 | UK | 2004 | starling | 26 | 43 | 9 | 101 | 8 | 46 | 21 | 1020 |
| WB CW04604 c11.6.04 | UK | 2004 | starling | 26 | 43 | 9 | 101 | 8 | 46 | 21 | 1020 |
| WB CW04607 c11.6.04 | UK | 2004 | starling | 26 | 43 | 9 | 101 | 8 | 46 | 21 | 1020 |
| WB CW04608 c11.6.04 | UK | 2004 | starling | 26 | 43 | 9 | 101 | 8 | 46 | 21 | 1020 |
| WB CW04610 c11.6.04 | UK | 2004 | starling | 26 | 43 | 9 | 101 | 8 | 46 | 21 | 1020 |
| WB CW04612 c11.6.04 | UK | 2004 | starling | 26 | 43 | 9 | 101 | 8 | 46 | 21 | 1020 |
| WB CW04613 c11.6.04 | UK | 2004 | starling | 26 | 43 | 9 | 101 | 8 | 46 | 21 | 1020 |
| WB CW04614 c11.6.04 | UK | 2004 | starling | 35 | 2 | 8 | 51 | 121 | 46 | 21 | 818 |
| WB CT86761 c11.6.04 | UK | 2004 | starling | 26 | 43 | 9 | 101 | 8 | 46 | 21 | 1020 |
| WB CT86879 c11.6.04 | UK | 2004 | starling | 35 | 2 | 8 | 51 | 121 | 46 | 21 | 818 |
| WB CT86880 c11.6.04 | UK | 2004 | starling | 35 | 2 | 9 | 51 | 8 | 2 | 21 | 1027 |
| WB GR/P c15.6.04 | UK | 2004 | starling | 35 | 2 | 8 | 51 | 121 | 46 | 21 | 818 |
| WB CW04616 c15.6.04 | UK | 2004 | starling | 35 | 2 | 8 | 51 | 121 | 46 | 21 | 818 |
| WB CW04626 c15.6.04 | UK | 2004 | starling | 26 | 43 | 9 | 101 | 8 | 46 | 21 | 1020 |
| WB CW04628 c15.6.04 | UK | 2004 | starling | 26 | 43 | 9 | 101 | 8 | 46 | 21 | 1020 |
| WB CW04630 c15.6.04 | UK | 2004 | starling | 26 | 43 | 9 | 101 | 8 | 46 | 21 | 1020 |
| WB CW04636 c15.6.04 | UK | 2004 | starling | 17 | 2 | 9 | 51 | 121 | 46 | 21 | 686 |
| WB CW04638 c15.6.04 | UK | 2004 | starling | 26 | 43 | 9 | 101 | 8 | 46 | 21 | 1020 |
| WB CW04639 c15.6.04 | UK | 2004 | starling | 35 | 2 | 8 | 51 | 121 | 46 | 21 | 818 |
| WB CW04644 c15.6.04 | UK | 2004 | starling | 26 | 43 | 9 | 101 | 8 | 46 | 21 | 1020 |

| | | | | | | | | | | | |
|---|---|---|---|---|---|---|---|---|---|---|---|
| WB CW04646 c15.6.04 | UK | 2004 | starling | 17 | 2 | 9 | 51 | 121 | 46 | 21 | 686 |
| WB CW04648 c15.6.04 | UK | 2004 | starling | 26 | 43 | 9 | 101 | 8 | 46 | 21 | 1020 |
| WB CW04651 c15.6.04 | UK | 2004 | starling | 17 | 2 | 9 | 51 | 121 | 46 | 21 | 686 |
| WB CW04652 c15.6.04 | UK | 2004 | starling | 35 | 2 | 8 | 51 | 121 | 46 | 21 | 818 |
| WB CW04653 c15.6.04 | UK | 2004 | starling | 17 | 2 | 9 | 51 | 121 | 46 | 21 | 686 |
| WB CT43821 c15.6.04 | UK | 2004 | starling | 17 | 2 | 9 | 51 | 121 | 46 | 21 | 686 |
| WB CT86804 c15.6.04 | UK | 2004 | starling | 26 | 43 | 9 | 101 | 8 | 46 | 21 | 1020 |
| WB CT86858 c15.6.04 | UK | 2004 | starling | 26 | 43 | 9 | 101 | 8 | 46 | 21 | 1020 |
| WB CW04659 c16.6.04 | UK | 2004 | starling | 26 | 43 | 9 | 101 | 8 | 46 | 21 | 1020 |
| WB CW04662 c16.6.04 | UK | 2004 | starling | 26 | 43 | 9 | 101 | 8 | 46 | 21 | 1020 |
| WB CW04663 c16.6.04 | UK | 2004 | starling | 26 | 43 | 9 | 5 | 121 | 46 | 21 | 1503 |
| WB CW04666 c16.6.04 | UK | 2004 | starling | 26 | 43 | 9 | 101 | 8 | 46 | 21 | 1020 |
| WB CT86780 c16.6.04 | UK | 2004 | starling | 26 | 43 | 9 | 101 | 8 | 46 | 21 | 1020 |
| WB CT86819 c16.6.04 | UK | 2004 | starling | 17 | 2 | 9 | 51 | 121 | 46 | 21 | 686 |
| WB CT86835 c16.6.04 | UK | 2004 | starling | 26 | 43 | 9 | 101 | 8 | 46 | 21 | 1020 |
| WB CT86852 c16.6.04 | UK | 2004 | starling | 26 | 43 | 9 | 101 | 8 | 46 | 21 | 1020 |
| WB CW04676 c18.6.04 | UK | 2004 | starling | 26 | 43 | 9 | 101 | 8 | 46 | 21 | 1020 |
| WB CW04678 c18.6.04 | UK | 2004 | starling | 35 | 2 | 8 | 51 | 121 | 46 | 21 | 818 |
| WB CW04679 c18.6.04 | UK | 2004 | starling | 26 | 43 | 9 | 101 | 8 | 46 | 21 | 1020 |
| WB CW04681 c18.6.04 | UK | 2004 | starling | 17 | 2 | 9 | 51 | 121 | 46 | 21 | 686 |
| WB CW04682 c18.6.04 | UK | 2004 | starling | 17 | 2 | 9 | 5 | 8 | 46 | 21 | 1021 |
| WB CW04683 c18.6.04 | UK | 2004 | starling | 17 | 2 | 9 | 51 | 121 | 46 | 21 | 686 |
| WB CW04684 c18.6.04 | UK | 2004 | starling | 35 | 2 | 9 | 51 | 8 | 2 | 21 | 1027 |
| WB CW04686 c18.6.04 | UK | 2004 | starling | 17 | 2 | 9 | 51 | 121 | 46 | 21 | 686 |
| WB CT86780 c18.6.04 | UK | 2004 | starling | 26 | 43 | 9 | 101 | 8 | 46 | 21 | 1020 |
| WB CT86851 c18.6.04 | UK | 2004 | starling | 17 | 2 | 9 | 5 | 8 | 46 | 21 | 1021 |
| WB CT86880 c18.6.04 | UK | 2004 | starling | 26 | 43 | 9 | 101 | 8 | 46 | 21 | 1020 |
| WB CW04602 c18.6.04 | UK | 2004 | starling | 26 | 43 | 9 | 101 | 8 | 46 | 21 | 1020 |
| WB CW04656 c18.6.04 | UK | 2004 | starling | 26 | 43 | 9 | 101 | 8 | 46 | 21 | 1020 |
| WB CW04688 c22.6.04 | UK | 2004 | starling | 35 | 2 | 9 | 51 | 8 | 2 | 21 | 1027 |
| WB CW04689 c22.6.04 | UK | 2004 | starling | 26 | 43 | 9 | 101 | 8 | 46 | 21 | 1020 |
| WB CW04692 c22.6.04 | UK | 2004 | starling | 26 | 2 | 9 | 51 | 121 | 46 | 21 | 1542 |
| WB CW04693 c22.6.04 | UK | 2004 | starling | 35 | 2 | 9 | 51 | 8 | 2 | 21 | 1027 |
| WB CW04696 c22.6.04 | UK | 2004 | starling | 26 | 43 | 9 | 101 | 8 | 46 | 21 | 1020 |
| WB CW04697 c22.6.04 | UK | 2004 | starling | 35 | 2 | 8 | 51 | 121 | 46 | 21 | 818 |
| WB CW04698 c22.6.04 | UK | 2004 | starling | 26 | 43 | 9 | 101 | 8 | 46 | 21 | 1020 |
| WB CW04701 c22.6.04 | UK | 2004 | starling | 26 | 43 | 9 | 101 | 8 | 46 | 21 | 1020 |
| WB CW04702 c22.6.04 | UK | 2004 | starling | 26 | 43 | 9 | 101 | 8 | 46 | 21 | 1020 |

| | | | | | | | | | | | |
|---|---|---|---|---|---|---|---|---|---|---|---|
| WB CW04706 c22.6.04 | UK | 2004 | starling | 26 | 43 | 9 | 101 | 8 | 46 | 21 | 1020 |
| CW04772 c24.11.04 | UK | 2004 | starling | 26 | 43 | 9 | 51 | 121 | 46 | 21 | 1022 |
| CW04774 c24.11.04 | UK | 2004 | starling | 26 | 43 | 9 | 51 | 121 | 46 | 21 | 1022 |
| CW04776 c24.11.04 | UK | 2004 | starling | 26 | 43 | 9 | 51 | 121 | 46 | 21 | 1022 |
| CW04780 c24.11.04 | UK | 2004 | starling | 26 | 43 | 9 | 51 | 121 | 46 | 21 | 1022 |
| CW04785 c26.11.04 | UK | 2004 | starling | 26 | 43 | 9 | 51 | 121 | 46 | 21 | 1022 |
| CW04789 c26.11.04 | UK | 2004 | starling | 26 | 43 | 9 | 51 | 121 | 46 | 21 | 1022 |
| CW04791 c30.11.04 | UK | 2004 | starling | 26 | 43 | 9 | 51 | 121 | 46 | 21 | 1022 |
| CW10311 c2.12.04 | UK | 2004 | starling | 35 | 2 | 9 | 51 | 8 | 2 | 21 | 1027 |
| CW04772 c26.11.04 | UK | 2004 | starling | 26 | 43 | 9 | 51 | 121 | 46 | 21 | 1022 |
| CW04779 c26.11.04 | UK | 2004 | starling | 26 | 43 | 9 | 51 | 121 | 46 | 21 | 1022 |
| CL95078 c30.11.04 | UK | 2004 | starling | 26 | 43 | 9 | 51 | 121 | 46 | 21 | 1022 |
| CL95078 c1.12.04 | UK | 2004 | starling | 26 | 43 | 9 | 51 | 121 | 46 | 21 | 1022 |
| CW04790 c9.12.04 | UK | 2004 | starling | 26 | 43 | 9 | 51 | 121 | 46 | 21 | 1022 |
| CW04605 c11.6.04 | UK | 2004 | starling | 26 | 43 | 9 | 101 | 8 | 46 | 21 | 1020 |
| CW04609 c11.6.04 | UK | 2004 | starling | 26 | 2 | 9 | 51 | 8 | 46 | 21 | 682 |
| WB CW04683 c22.6.04 | UK | 2004 | starling | 35 | 2 | 9 | 51 | 8 | 2 | 21 | 1027 |
| WB CW04710 c24.6.04 | UK | 2004 | starling | 17 | 2 | 9 | 51 | 121 | 46 | 21 | 686 |
| WB CW04712 c24.6.04 | UK | 2004 | starling | 26 | 43 | 9 | 101 | 8 | 46 | 21 | 1020 |
| WB CW04714 c24.6.04 | UK | 2004 | starling | 26 | 43 | 9 | 101 | 8 | 46 | 21 | 1020 |
| WB CW04716 c24.6.04 | UK | 2004 | starling | 26 | 43 | 9 | 101 | 8 | 46 | 21 | 1020 |
| WB CW04718 c24.6.04 | UK | 2004 | starling | 35 | 2 | 8 | 51 | 121 | 46 | 21 | 818 |
| WB CW04721 c24.6.04 | UK | 2004 | starling | 26 | 43 | 9 | 101 | 8 | 46 | 21 | 1020 |
| WB CW04722 c24.6.04 | UK | 2004 | starling | 17 | 2 | 9 | 5 | 8 | 46 | 21 | 1021 |
| WB CW04723 c24.6.04 | UK | 2004 | starling | 26 | 43 | 9 | 101 | 8 | 46 | 21 | 1020 |
| WB CW04728 c24.6.04 | UK | 2004 | starling | 17 | 2 | 9 | 51 | 121 | 46 | 21 | 686 |
| WB CW04696 c24.6.04 | UK | 2004 | starling | 26 | 43 | 9 | 101 | 8 | 46 | 21 | 1020 |
| WB CW04731 c25.6.04 | UK | 2004 | starling | 35 | 2 | 8 | 51 | 121 | 46 | 21 | 818 |
| WB CW04735 c25.6.04 | UK | 2004 | starling | 26 | 43 | 9 | 101 | 8 | 46 | 21 | 1020 |
| WB CW04736 c25.6.04 | UK | 2004 | starling | 26 | 43 | 9 | 101 | 8 | 46 | 21 | 1020 |
| WB CT86776 c25.6.04 | UK | 2004 | starling | 17 | 2 | 9 | 51 | 121 | 46 | 21 | 686 |
| WB CT86897 c25.6.04 | UK | 2004 | starling | 17 | 2 | 9 | 51 | 121 | 46 | 21 | 686 |
| WB CW04605 c25.6.04 | UK | 2004 | starling | 26 | 2 | 9 | 5 | 8 | 46 | 21 | 1392 |
| WB CW04609 c25.6.04 | UK | 2004 | starling | 17 | 2 | 9 | 51 | 121 | 46 | 21 | 686 |
| WB CW04658 c25.6.04 | UK | 2004 | starling | 35 | 2 | 9 | 51 | 8 | 2 | 21 | 1027 |
| WB CW04712 c25.6.04 | UK | 2004 | starling | 26 | 43 | 9 | 101 | 8 | 46 | 21 | 1020 |
| WB CW04713 c25.6.04 | UK | 2004 | starling | 35 | 2 | 9 | 51 | 8 | 2 | 21 | 1027 |
| WB CW04714 c25.6.04 | UK | 2004 | starling | 26 | 43 | 9 | 101 | 8 | 46 | 21 | 1020 |

| | | | | | | | | | | | |
|---|---|---|---|---|---|---|---|---|---|---|---|
| WB CW04704 c29.6.04 | UK | 2004 | starling | 26 | 43 | 9 | 101 | 8 | 46 | 21 | 1020 |
| WB CW04714 c29.6.04 | UK | 2004 | starling | 35 | 2 | 9 | 51 | 8 | 2 | 21 | 1027 |
| WB CW04683 c30.6.04 | UK | 2004 | starling | 35 | 2 | 9 | 51 | 8 | 2 | 21 | 1027 |
| 007A-0838 | Canada: Quebec | 2006 | environmental water | 26 | 2 | 9 | 51 | 8 | 46 | 1 | 4203 |
| PIK3 | Finland | 2005 | Unk. | 17 | 2 | 8 | 51 | 8 | 46 | 21 | 4573 |
| VDL7103 | USA: Iowa | 2008 | sheep | 35 | 2 | 9 | 432 | 8 | 46 | 21 | 5128 |
| MIDO 818 (5) | USA: Georgia | 2005 | environmental water | 35 | 2 | 9 | 51 | 8 | 2 | 21 | 1027 |
| NORO 516 (12) | USA: Georgia | 2005 | environmental water | 26 | 2 | 9 | 51 | 8 | 46 | 21 | 682 |
| MIDO 301 (13) | USA: Georgia | 2005 | environmental water | 26 | 2 | 9 | 51 | 8 | 46 | 21 | 682 |
| MIDO 812 (78) | USA: Georgia | 2005 | environmental water | 35 | 2 | 9 | 51 | 8 | 2 | 21 | 1027 |
| CSSS_66821 | UK: Nottinghamshire | 2002 | human: unspecified | 26 | 2 | 9 | 51 | 8 | 46 | 21 | 682 |
| 007A-0377 | Canada: Quebec | 2005 | environmental water | 35 | 2 | 9 | 51 | 8 | 2 | 21 | 1027 |
| 007A-0385 | Canada: Quebec | 2005 | environmental water | 26 | 2 | 9 | 51 | 8 | 46 | 21 | 682 |
| 007A-0391 | Canada: Quebec | 2005 | environmental water | 26 | 2 | 9 | 51 | 8 | 46 | 21 | 682 |
| 007A-0792 | Canada: Quebec | 2006 | environmental water | 26 | 2 | 9 | 51 | 8 | 46 | 21 | 682 |
| 007A-0828 | Canada: Quebec | 2006 | environmental water | 26 | 2 | 9 | 51 | 8 | 46 | 21 | 682 |
| 06_BB_25 | Australia | 2006 | blackbird | 35 | 2 | 9 | 51 | 8 | 46 | 21 | 3068 |
| jonas_star_3 | Sweden | 2000 | starling | 26 | 2 | 9 | 51 | 8 | 46 | 21 | 682 |
| jonas_star_7 | Sweden | 2000 | starling | 26 | 2 | 9 | 51 | 8 | 46 | 21 | 682 |
| jonas_star_5 | Sweden | 2000 | starling | 26 | 2 | 9 | 51 | 8 | 46 | 21 | 682 |
| jonas_star_1 | Sweden | 2000 | starling | 26 | 2 | 9 | 51 | 8 | 46 | 21 | 682 |
| 2011D-8859 | USA | Unk. | Unk. | 35 | 2 | 9 | 534 | 8 | 46 | 21 | 6677 |
| J-Br-706 | USA: Ohio | 2010 | starling | 26 | 43 | 9 | 101 | 8 | 46 | 21 | 1020 |
| E-Br-2 | USA: Ohio | 2010 | starling | 26 | 43 | 9 | 101 | 8 | 46 | 21 | 1020 |
| I-Br-15 | USA: Ohio | 2010 | starling | 26 | 43 | 9 | 101 | 8 | 46 | 21 | 1020 |
| I-Br-14 | USA: Ohio | 2010 | starling | 26 | 43 | 9 | 101 | 8 | 46 | 21 | 1020 |
| H-Br-7 | USA: Ohio | 2010 | starling | 17 | 2 | 9 | 5 | 8 | 46 | 21 | 1021 |
| LB_BS3.1_21C2 | Thailand: Bangkok | 2012 | broiler environment | 35 | 2 | 9 | 5 | 8 | 553 | 21 | 6995 |
| E120455 | Luxembourg | 2012 | environmental water | 17 | 2 | 9 | 5 | 8 | 46 | 21 | 1021 |
| W0004 | UK: Scotland | 2015 | wild bird | 26 | 2 | 9 | 51 | 8 | 46 | 21 | 682 |
| BS3.121C2 | Thailand | 2012 | broiler environment | 35 | 2 | 9 | 5 | 8 | 553 | 21 | 6995 |
| B1432b | New Zealand: Manawatu | 2009 | wild bird | 35 | 43 | 9 | 5 | 8 | 46 | 21 | 681 |
| B1624b | New Zealand: Manawatu | 2009 | wild bird | 26 | 2 | 9 | 51 | 8 | 46 | 5 | 208 |
| W860b | New Zealand: Waikato | 2013 | environmental water | 26 | 2 | 9 | 51 | 8 | 46 | 21 | 682 |
| CL95044_4.6.4 | UK | Unk. | starling | 26 | 43 | 9 | 101 | 8 | 46 | 21 | 1020 |
| starling1020 | UK | Unk. | starling | 26 | 43 | 9 | 101 | 8 | 46 | 21 | 1020 |
| SGEHI2013-C591-1 | Singapore | 2013 | wild bird | 35 | 43 | 8 | 51 | 8 | 46 | 21 | 9603 |
| PNUSAC005563 | USA | Unk. | Unk. | 26 | 2 | 9 | 51 | 8 | 46 | 21 | 682 |

| | | | | | | | | | | | |
|---|---|---|---|---|---|---|---|---|---|---|---|
| PNUSAC005518 | USA | Unk. | Unk. | 26 | 2 | 9 | 51 | 8 | 46 | 21 | 682 |
| FSIS11920083 | USA: California | 2019 | chicken offal or meat | 35 | 2 | 9 | 432 | 8 | 46 | 21 | 5128 |
| FSIS1607521 | USA: California | 2016 | cattle | 35 | 2 | 8 | 51 | 121 | 46 | 21 | 818 |
| FSIS11813822 | USA: Minnesota | 2018 | cattle | 26 | 2 | 9 | 51 | 8 | 46 | 21 | 682 |
| **SKBC94** | **USA: North Carolina** | **2016** | **black bear** | 26 | 2 | 9 | 51 | 8 | 46 | 21 | 682 |
| 125 | UK | Unk. | starling | 26 | 43 | 9 | 101 | 8 | 46 | 21 | 1020 |
| 10352 | USA | Unk. | starling | 35 | 2 | 281 | 5 | 8 | 46 | 21 | 12448 |
| CMB210274 | New Zealand | 2021 | environmental water | 237 | 2 | 9 | 5 | 8 | 222 | 21 | 11295 |
| CMB210352 | New Zealand | 2021 | environmental water | 237 | 2 | 9 | 5 | 8 | 222 | 21 | 11295 |
| B203-270820-01 | Luxembourg | 2020 | wild bird | 4 | 43 | 9 | 5 | 8 | 46 | 21 | 11385 |
| FSIS12106873 | USA | 2021 | calf | 26 | 2 | 9 | 51 | 8 | 46 | 21 | 682 |
| OXCBB-3596 | UK | 2004 | chicken | 26 | 43 | 9 | 51 | 121 | 46 | 21 | 1022 |
| OXCBB-3684 | UK | 2004 | chicken | 26 | 43 | 9 | 51 | 121 | 46 | 21 | 1022 |
| OXCBB-3577 | UK | 2004 | chicken | 26 | 43 | 9 | 51 | 121 | 46 | 21 | 1022 |
| OXCBB-3478 | UK | 2004 | chicken | 26 | 43 | 9 | 51 | 121 | 46 | 21 | 1022 |
| OXCBB-3510 | UK | 2004 | chicken | 26 | 43 | 9 | 51 | 121 | 46 | 21 | 1022 |
| OXCstar152 | UK | 2004 | chicken | 26 | 43 | 9 | 51 | 121 | 46 | 21 | 1022 |
| W69 | Thailand | 2018 | environmental water | 26 | 2 | 9 | 51 | 8 | 46 | 21 | 682 |

**Supplementary Table 4: Strains in PubMLST that share alleles with either SKBC3 or SKBC5**

| Strain | Prov/State/Region | Country | Year | Month | disease | source |
|---|---|---|---|---|---|---|
| 007A-0615 | Quebec | Canada | 2006 | | | environmental waters |
| VDL26018 | Iowa | USA | 2011 | | gastroenteritis | dog |
| SFBRC-47 | Georgia | USA | 2013 | 11 | | environmental waters |
| SFBRC-61 | Georgia | USA | 2013 | 12 | | environmental waters |
| PNUSAC007623 | NA | USA | NA | | NA | NA |
| PNUSAC006289 | HHS Region 5 | USA | 2018 | | | human |
| PNUSAC006563 | HHS Region 8 | USA | 2018 | | | human |
| SKBC3 | North Carolina | USA | 2014 | 7 | carrier | black bear |
| SKBC5 | North Carolina | USA | 2014 | 8 | carrier | black bear |
| SFBRC-36 | Georgia | USA | 2013 | 11 | | environmental waters |
| SFBRC-59 | Georgia | USA | 2013 | 12 | | environmental waters |
| SFBRC-60 | Georgia | USA | 2013 | 12 | | environmental waters |
| SFBRC-66 | Georgia | USA | 2013 | 12 | | environmental waters |

Alleles from SKBC3 or SKBC5 highlighted in orange are present in more than one strain, while those highlig

| epidemiology | species | aspA | glnA | gltA | glyA | pgm | tkt | uncA | ST |
|---|---|---|---|---|---|---|---|---|---|
| environmental isolate | Campylobacter jejuni | 255 | 339 | 279 | 383 | 479 | 391 | 276 | 4361 |
| | Campylobacter jejuni | 305 | 6 | 61 | 176 | 40 | 180 | 3 | 6971 |
| environmental isolate | Campylobacter jejuni | 305 | 409 | 339 | 664 | 578 | 391 | 344 | 7945 |
| environmental isolate | Campylobacter jejuni | 305 | 6 | 61 | 176 | 40 | 180 | 224 | 7943 |
| NA | Campylobacter jejuni | 305 | 6 | 137 | 176 | 40 | 32 | 3 | 10539 |
| | Campylobacter jejuni | 305 | 6 | 61 | 176 | 40 | 180 | 3 | 6971 |
| | Campylobacter jejuni | 305 | 6 | 61 | 176 | 40 | 180 | 3 | 6971 |
| carrier | Campylobacter jejuni | 255 | 530 | 279 | 607 | 740 | 585 | 276 | 7630 |
| carrier | Campylobacter jejuni | 305 | 756 | 279 | 607 | 479 | 585 | 276 | 10620 |
| environmental isolate | Campylobacter jejuni | 305 | 6 | 61 | 176 | 40 | 180 | 224 | 7943 |
| environmental isolate | Campylobacter jejuni | 305 | 6 | 61 | 176 | 40 | 180 | 224 | 7943 |
| environmental isolate | Campylobacter jejuni | 305 | 6 | 61 | 176 | 40 | 180 | 224 | 7943 |
| environmental isolate | Campylobacter jejuni | 426 | 530 | 339 | 607 | 779 | 391 | 344 | 7949 |

hte in yellow were present in only the particular isolate.

**clonal complex**

ST-179 complex

**Supplementary Table 5: Similarity to subset of RM1221 CJIE1 (Mu-like) genes.**

| | | **Strain** (Location) | | | | | |
|---|---|---|---|---|---|---|---|
| **RM1221 locus tags** | **Description** | **SKBC1** (near *npdA*) | **SKBC3-1** (near *rarA*) | **SKBC3-2** (near *ctsT*) | **SKBC5-1** (in CJIE2-like) | **SKBC5-2** (in CJIE3-like) | **SKBC25** (near *selU*) |
| CJE0270 | DNA transposition protein A | **44.0%** | **90.9%** | **44.7%** | **90.1%** | **43.7%** | **86.2%** |
| CJE0269 | DNA transposition protein B | **46.9%** | **95.7%** | **46.4%** | **95.7%** | **46.4%** | **95.8%** |
| CJE0265 | host-nuclease inhibitor protein Gam | **92.4%** | **89.6%** | **94.0%** | **89.6%** | **94.0%** | **90.9%** |
| CJE0256 | *dns* (extracellular DNase) | **-** | **92.9%** | **-** | **92.9%** | **-** | **99.0%** |
| CJE0254 | tail D protein | **57.9%** | **96.6%** | **58.1%** | **96.6%** | **58.1%** | **97.6%** |
| CJE0252 | tail protein | **70.0%** | **96.8%** | **69.9%** | **96.8%** | **69.9%** | **99.7%** |
| CJE0251 | Mu-like prohage F protein | **-** | **95.7%** | **-** | **95.7%** | **-** | **99.6%** |
| CJE0244 | Mu-like prohage I protein | **-** | **95.7%** | **-** | **95.7%** | **-** | **99.6%** |
| CJE0236 | baseplate assembly protein V | **97.4%** | **97.4%** | **97.4%** | **97.2%** | **97.4%** | **97.9%** |
| CJE0235 | baseplate assembly protein W | **98.6%** | **92.1%** | **92.1%** | **92.1%** | **90.7%** | **97.9%** |
| CJE0233 | baseplate assembly protein J | **97.1%** | **94.7%** | **94.7%** | **94.7%** | **94.9%** | **96.1%** |
| CJE0232 | tail protein | **96.9%** | **96.8%** | **94.2%** | **96.8%** | **95.8%** | **96.8%** |
| CJE0231 | tail fiber H protein | **94.2%** | **96.8%** | **96.8%** | **96.8%** | **96.9%** | **95.8%** |
| CJE0227 | major tail sheath protein | **97.8%** | **95.4%** | **95.5%** | **95.7%** | **95.8%** | **97.9%** |
| CJE0226 | major tail tube protein | **71.5%** | **96.5%** | **71.5%** | **96.7%** | **71.5%** | **96.5%** |
| CJE0222 | tail tape measure protein | **42.40%** | **77.4%** | **42.60%** | **77.4%** | **42.60%** | **77.6%** |
